# Discovery and Development of DC-174 as a Novel Oral Snakebite Treatment

**DOI:** 10.1101/2025.05.23.655830

**Authors:** Daniel J.W. Chong, Laura-Oana Albulescu, Adam P. Westhorpe, Rachel H. Clare, Amy E. Marriott, Christopher M. Woodley, Ramachandran Gunasekar, Nada Mosallam, Edouard Crittenden, Emma Stars, Charlotte A. Dawson, Jeroen Kool, Mark C. Wilkinson, Suet C. Leung, Neil G. Berry, Nicholas R. Casewell, Paul M. O’Neill

**Affiliations:** Department of Chemistry, University of Liverpool, Grove Street, Liverpool, L69 7ZD, UK; Centre for Snakebite Research & Interventions, Liverpool School of Tropical Medicine, Pembroke Place, Liverpool, L3 5QA, UK; Department of Chemistry and Pharmaceutical Sciences, Amsterdam Institute for Molecular and Life Sciences, Vrije Universiteit Amsterdam, De Boelelaan 1105, Amsterdam 1081 HV, the Netherlands

## Abstract

Snakebite envenoming is a neglected tropical disease that causes high mortality and morbidity. The current treatment, intravenous antivenom, comes with numerous disadvantages making new therapeutics important. Optimised small molecules offer the possibility for oral use at the onset of envenoming, and the highly pathogenic, zinc-dependent, snake venom metalloproteinase toxin family represents an attractive target for drug discovery. Through systematic chemical modification guided by molecular modelling, we describe the development of hydroxamic acid DC-174, a molecule that displays potent broad spectrum metalloproteinase inhibition and neutralises the procoagulant activities of multiple snake venoms. In oral-dosing studies, DC-174 showed preclinical efficacy in a mouse model of severe envenoming, with efficacy boosted by a pharmacokinetically-informed multiple dosing regimen. This rationally designed metalloproteinase inhibitor offers a potential paradigm shift from delayed treatment with antivenom in tertiary hospitals to a contemporary approach using oral drugs amenable for rapid use in snakebite-affected communities.

## Background

Snakebite envenoming is a life-threatening condition predominantly affecting people in rural communities in Africa, Asia and Latin America, and results in an annual death toll of up to 138,000, with many more survivors suffering long term morbidity^1^. In 2017 the World Health Organization (WHO) reclassified snakebite envenoming as a priority Neglected Tropical Disease^2^ and a roadmap was developed to reduce the snakebite burden, with one key pillar of this strategy being to ensure access to safe and effective treatment, inclusive of improved therapeutics^3^.

Antivenoms are current standard of care snakebite treatments that consist of animal-derived polyclonal antibodies sourced from hyper-immunised animals. Despite saving lives, antivenoms have several limitations, including the requirement for administration in a clinical environment due to the necessity for intravenous delivery, management of adverse reactions and reliance on cold chain storage^4^. Placing these constraints on patients who are often on average six hours from a hospital setting^5,6^, likely contributes to poor outcomes, given the often acute and life-threatening nature of snakebite. Additional challenges include the low affordability and accessibility of antivenom in many tropical regions, which is partly due to the necessity to manufacture different products for different regions, due to the complex and variable nature of venom toxins among snake species^7^.

Despite extensive venom variation, four main pathogenic toxin families have been identified as being of greatest importance for snakebite due to their relative abundance and toxicity; the snake venom metalloproteinases (SVMPs), snake venom serine proteases (SVSPs), phospholipases A_2_ (PLA_2_s) and three-finger toxins (3FTxs)^8^. The SVMPs are zinc-dependent enzymes present in the venom of most snakes, but are particularly abundant in viper venoms (*Viperidae*)^8,9^. SVMPs are divided into three subclasses (P-I, P-II, P-III) based on gene organisation; though each contains structurally and functionally distinct toxin isoforms^9^. SVMPs induce and prolong local and systemic haemorrhage by degrading extracellular matrix proteins resulting in microvascular damage and by cleaving blood clotting factors, such as fibrinogen and prothrombin, resulting in coagulopathy^10,11^. Their high abundance in many viper venoms (up to 74% of all toxins^8^) makes SVMPs a highly attractive target for novel snakebite therapeutics.

One contemporary approach (among others^12^) recently applied to more broadly inhibit diverse toxin isoforms found across snake species has been the use of small molecule drugs^13–17^. Such molecules possess several theoretical advantages over current antivenom, including improved cross-snake species neutralisation, increased affordability and no requirement for cold-chain, but also offer the possibility of oral formulation for rapid administration in the community after a snakebite, thus reducing the critical time to treatment^18^. Recently, several drugs that may generically target the active site of venom enzymes have been actively repurposed for snakebite, including the PLA_2_ inhibitor varespladib, which showed impressive preclinical efficacy and recently completed a Phase II snakebite trial, though the primary endpoint was not met^19,20^. Similarly, two distinct group of molecules have been reported to show promising SVMP-inhibitory activities. Metal chelating molecules, particularly 2,3-dimercapto-1- propanesulfonic acid (DMPS), showed preclinical protection in *in vivo* envenoming models^14,16^ and has since been used in a Phase I dose optimisation study to support onward clinical development for snakebite indication^21^. The second comprises several matrix metalloproteinase (MMP) inhibitors. These compounds feature a zinc-binding motif which directly engages with the active site of SVMPs^22^, and include marimastat^15,16,23–26^, prinomastat^24,26,27^ and batimastat^23–26^. These hydroxamic acids demonstrated potent inhibition of SVMP toxins *in vitro*, inhibition of procoagulant venom effects in physiologically relevant bioassays, and showed variable promise in preclinical trials for mitigating the local and systemic haemotoxic effects of snake venoms in small animal models^15,16,23,24^.

While such repurposing efforts have been highly encouraging, our understanding of the chemical space around SVMP inhibitors remains largely unexplored, as the identified repurposed drugs have not been optimised via medicinal chemistry approaches to improve their drugability for snakebite indication. To address this issue and expand the snakebite drug portfolio, we recently described a high-throughput screening (HTS) campaign that identified several potent SVMP inhibitors from a ∼3,500 compound library, including the MMP inhibitor XL-784, which displayed double digit nanomolar inhibition of the SVMP activity of several diverse snake venoms^26^. Using XL-784 as a starting scaffold, here we describe the first snakebite specific medicinal chemistry campaign to identify lead candidate SVMP inhibitors that display broad cross-snake species inhibitory activity, and with superior potency and drug metabolism and pharmacokinetic (DMPK) properties to current lead drugs. Following a rationally designed optimisation cascade, consisting of enzymatic and phenotypic assessments coupled to DMPK analyses, DC-174 was identified as a lead candidate molecule. This first, rationally designed, snakebite drug has desirable physicochemical and broad-spectrum venom inhibitory properties and demonstrates impressive preclinical efficacy against snakebite when dosed orally in a murine rescue model of envenoming.

## Results

### Optimisation of SVMP inhibitors identifies DC-174 as lead candidate

Our prior primary HTS campaign identified several strong hits, including the known MMP- inhibitors marimastat, prinomastat and XL-784^26^. XL-784 has a highly modifiable piperazine scaffold and consists of (1) primary scaffold, (2) 4-substituted aryl groups and (3) N1-side chain (Figure 1a). Based on previous structure-activity relationship (SAR) studies with human MMPs, we preserved the central core scaffold in our SAR optimisation strategy. In the related MMP-inhibitor Ro-1130830 (Figure 1b) and its analogues, the hydroxamic acid is shown to chelate the zinc while forming a hydrogen bond with a nearby glutamate residue^28^, with further hydrogen bonds predicted between the linking sulfonamide and backbone NH groups of a nearby loop^29,30^. The aromatic rings represent the S1’ binding substituent (Figure 1c) shown in the development of selective MMP13 inhibitors^28^. X-ray crystal structures of multiple SVMP toxins suggest this region and its available volume for binding may be variable (Supplementary figure 1), therefore modifications may affect the spectrum of SVMPs inhibited. Consequently, variation of substituents of the aromatic-ring and N1-side chain subunits were performed to investigate the structure-activity relationship against SVMPs.

**Figure 1.**
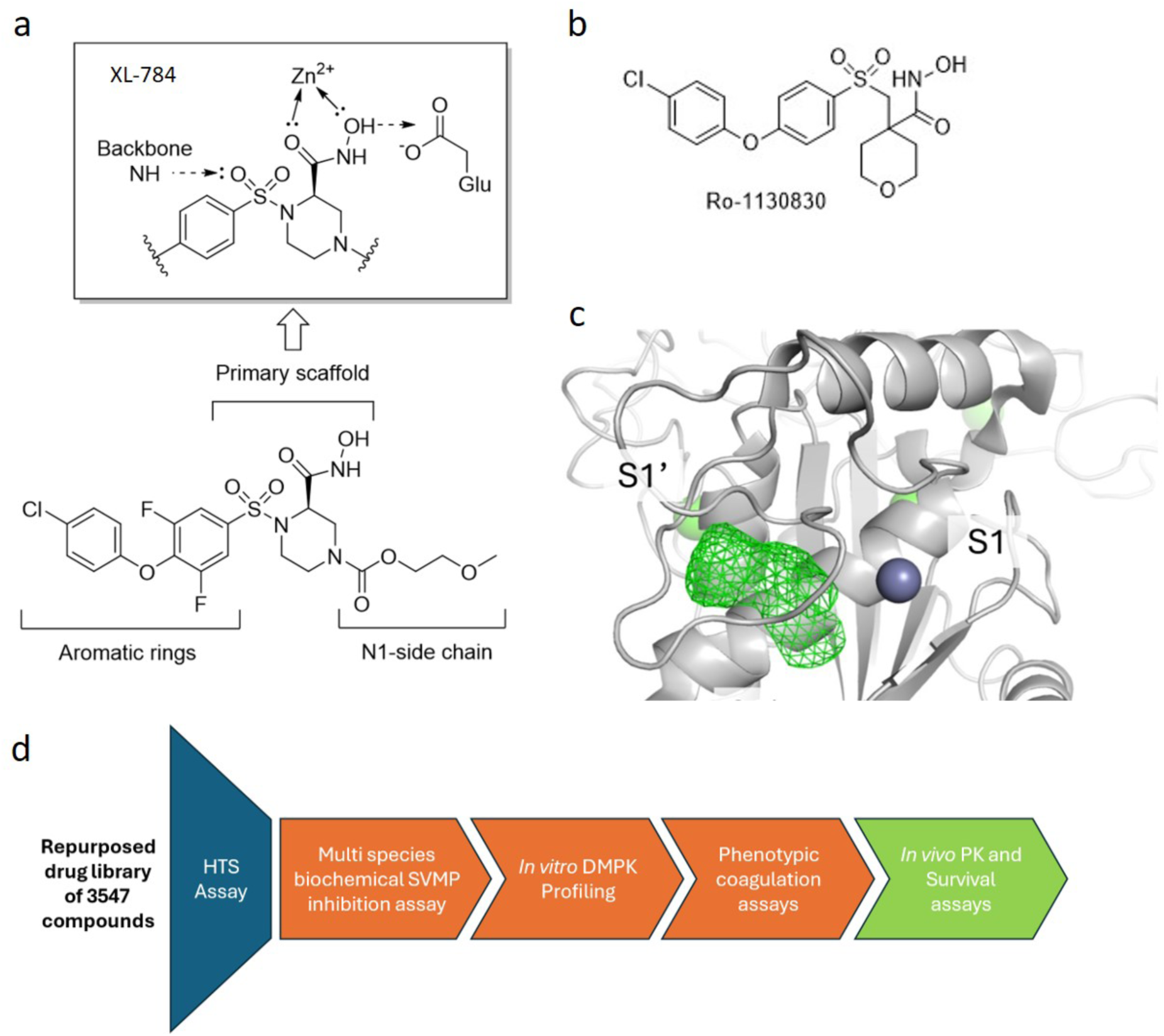
Medicinal chemistry optimisation strategy. a) Structure of the HTS hit compound XL-784 and proposed binding mode of the head group from studies using human MMP inhibitors. b) The chemical structure of MMP-inhibitor Ro-1130830. c) Representation of the binding site of the SVMP bothropasin from *Bothrops jararaca*. The protein secondary structure is rendered as grey ribbon, the catalytic zinc a as dark blue sphere and the S1’ pocket as green mesh with binding regions (S1 and S1’ pockets) labelled. d) The screening cascade for compounds produced in this study.

Throughout our medicinal chemistry campaign, compounds were evaluated for their ability to inhibit enzymatic SVMP toxin activity *in vitro* using a previously described fluorescent-based substrate cleavage assay^26^ against five medically important viper venoms, namely *Echis romani* (Roman’s carpet viper, West Africa), *Crotalus atrox* (Western diamondback rattlesnake, North America), *Calloselasma rhodostoma* (Malayan pit viper, South East Asia), *Bothrops jararaca* (jararaca, South America) and *Bitis arietans* (puff adder, Africa). We defined a potent inhibitor as having *in vitro* SVMP inhibition potency of less than 100 nM. The downstream testing cascade (Figure 1d) included assessment of *in vitro* DMPK properties of the compounds, with thresholds for “good” compounds defined as rat hepatocyte clearance <10 µL/min/10^6^ cells, human microsomal clearance <10 µL/min/mg intrinsic clearances, and aqueous solubility at pH 7.4 >500 µM^31,32^. Secondary inhibitory assays assessing neutralisation of venom activity on plasma clotting were used to triage synthesised molecules and benchmark against prinomastat and XL-784, before progression of lead compounds into *in vivo* models to evaluate pharmacokinetic (PK) and efficacy profiles.

The medicinal chemistry campaign resulted in 12 compounds (Figure 2, Supplementary table 1; chemical synthetic procedures can be found in the supplementary information). The para-substitution on the aromatic ring was initially accessed and hydroxamic acid **28** was able to retain similar potency to parent XL-784, while showing improvement in metabolic stability (rat hepatocyte clearance of 22.8 µl/min/10^6^ cells). Further SAR investigation of the ether linkage between the biphenyl motif showed that both para-halogenated **29** and **30** bolstered the overall SVMP potency profile, despite poor metabolic stability observed for the para-fluorinated **30**. The truncation of the biphenyl system demonstrated better aqueous solubility (>1000 µM) and significantly lower protein binding (82% free) for compound **20** (Supplementary table 1). Next, the methoxy ethyl side chain was replaced with a sulfonamide group which revealed effective inhibition of procoagulant venom activity by compounds **19** and **21-23** (X-ray crystallography analysis of **23** can be found in Supplementary table 2). This distinct secondary activity assay measured inhibition of a key parameter of clinical envenoming, coagulopathy^33^, which is often driven by SVMPs^11^. All sulfonamide *N*-1 analogues displayed measurable inhibition of procoagulant venom activity (Supplementary table 3), unlike the parent molecule XL-784, monoaryl **20**, and biaryl analogues **29** and **30**, which were all inactive in this assay. The thiomorpholine-based biaryl prinomastat also demonstrated inhibition of procoagulant activity, suggesting inactivity of **20**, **29** and **30** could be due to the *N*-1 substituted methoxyethyl group. This dependence on the *N*-1 substituent was highlighted by the observation that EC_50s_ increased with steric bulk of the *N*-1 substituent from a minimum at **23** (cyclopropyl), increasing with **22** (dimethyl-isoxazole) and **21** (methoxyphenyl). Overall, the compact cycloproyl sulfonamide **23** (named DC-174) emerged as the most desirable analogue in the series with improved broad-spectrum potent inhibition of SVMP activity (4.7 - 38.6 nM EC_50s_), metabolic stability (13.3 µl/min/10^6^ cells) and inhibition of procoagulant venom activity (13.4 and 135.0 nM EC_50s_ in *B. jararaca* and *E. romani*, respectively).

**Figure 2.**
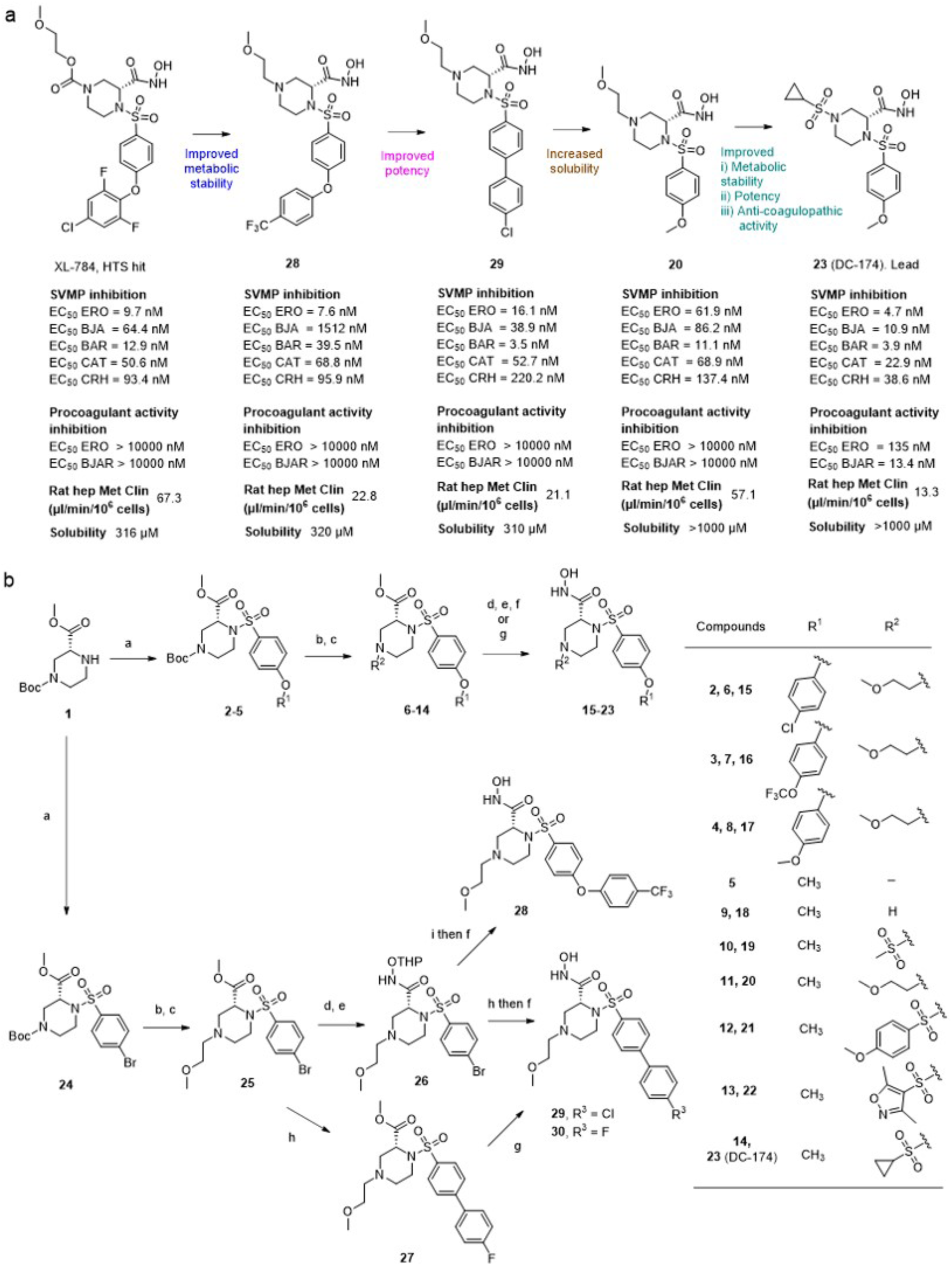
Hit-to-lead optimisation towards DC-174 (compound 23). a) The structural progression from XL-784 to DC-174. ERO, *E. romani*; BJA, *B. jararaca*; BAR, *Bitis arietans*; CAT, *Crotalus atrox*; CRH, *Calloselasma rhodostoma*. b) Chemical synthetic schemes for compounds **15**-**23** and **28**-**30**. Reaction condition: a) sulfonyl chlorides, NEt_3_, DMAP, 1,4- dioxane, rt, 16 h, 67-95%; b) TFA, DCM, rt, 16 h; c) Br(CH_2_)_2_OCH_3_, K_2_CO_3_, DMF, 60°C, 16 h, 32-71% or sulfonyl chlorides, NEt_3_, DMAP, 1,4-dioxane, rt, 16 h, 70-93%; d) NaOH, MeOH, rt, 1 h; e) EDC, HOBt, DMF, NMM, THPO-NH_2_, rt, 16 h, 51-95%; f) 4N HCl, 1,4-dioxane, rt, 1 h, 52-91%; g) NH_2_OH, KOCN, 1,4-dioxane, rt, 16 h, 47-94%; h) K_2_CO_3_, Pd(PPh_3_)_4_, 1,4-dioxane, H_2_O, 80°C, 79%; i) 4-(trifluoromethyl)phenol, K_3_PO_4_, Pd(OAc)_2_, di- tBuXPhos, toluene, 100°C, 16 h, 15%.

### DC-174 inhibits structurally diverse SVMPs and prevents venom-induced coagulation and haemorrhage

To explore whether DC-174 inhibits the breadth of structurally-distinct SVMP isoforms, we purified the P-I, P-II and P-III SVMPs from *E. romani* venom (Supplementary figure 2) and assessed their capacity to cleave the coagulation factors prothrombin and fibrinogen, before determining whether DC-174 could prevent this coagulotoxic activity. All three SVMP classes degraded fibrinogen and prothrombin to varying extents (Figure 3a), except for the P-II on prothrombin (Supplementary figure 3). Comparisons of DC-174 against parent molecule XL- 784 and prinomastat showed that all compounds protected against the degradation of fibrinogen, though XL-784 was inferior against P-II SVMP activity (Figure 3a, see lower molecular weight degradation bands). XL-784 was also unable to protect against prothrombin degradation by both P-I and P-III SVMPs, while DC-174 and prinomastat gave almost full protection, though DC-174 was superior based on visualisation of fewer degradation products (Figure 3b). These data could explain the differences in potency in the earlier coagulation assay, since prothrombin activation is a critical driver of procoagulant venom activity^11^.

**Figure 3.**
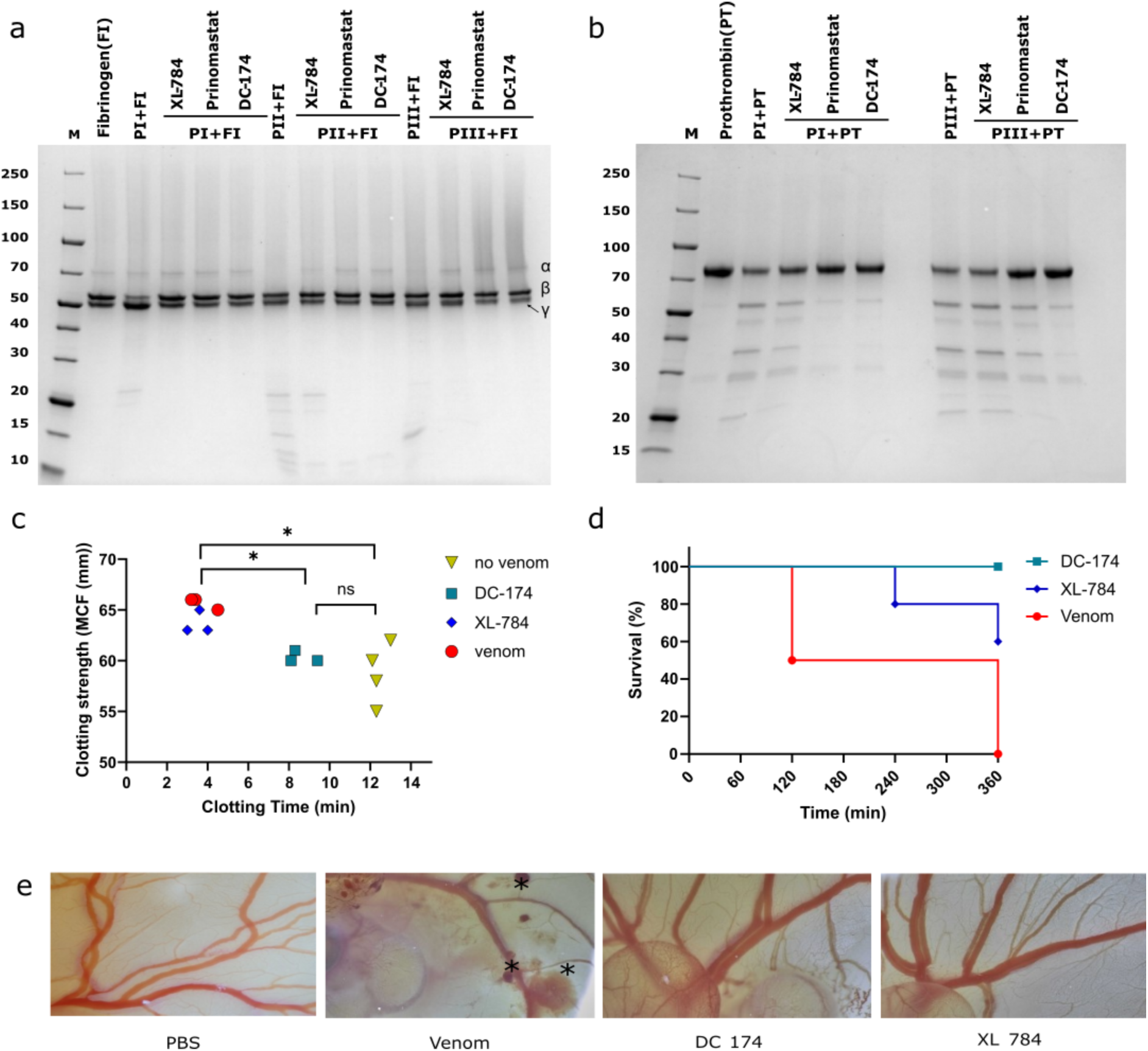
DC-174 reduces SVMP isoform-mediated cleavage of clotting factors and venom-induced coagulopathy and protects against lethality and haemorrhage in a chicken embryo model of envenoming. Inhibition of the degradation of a) fibrinogen and b) prothrombin by XL-784, prinomastat, and DC-174. The three different structural isoforms of *E. romani* SVMPs (P-I, P-II, P-III) were incubated for 30 mins in the presence of each drug after which fibrinogen or prothrombin were added, the samples incubated for 1 h, before visualisation of protein profiles via Coomassie staining of SDS-PAGE gel separations. The α, β and γ subunits of fibrinogen and indicated on the right side of the gel. c) Thromboelastographic measurements of clotting time and MCF resulting from human whole blood treated with *E. romani* venom (60 ng) co-administered with vehicle (DMSO - red circles), DC-174 (teal squares) and XL-784 (blue diamonds) compared with untreated controls (yellow triangles, n=4). Treatments were dosed at 5 µM (n=3). An ordinary one-way ANOVA was performed for all groups with significance indicated by asterisks (*, *p<*0.05; F(3,9) = 9.653, no venom vs XL-784, *p* = 0.029, venom vs. DC-174, *p* = 0.027). d) Survival curves of chicken embryos topically dosed with 10 µg of *E. romani* venom or venom immediately followed by 1 µg of either DC-174 or XL-784 (n=5). Embryos were observed at 1, 2, 4, and 6 hours post-exposure for the presence of an observable heartbeat. e) Representative images of the dosed embryos from (d) at 1-hour post-venom exposure. Images show *E. romani* venom only controls with extensive haemorrhage (asterisks), while those treated with DC-174 or XL- 784 displayed no evidence of haemorrhage at the same 1-hour timepoint. A PBS-dosed (healthy) negative control is shown for comparative purposes.

To further explore the improved venom inhibition of DC-174 over its parent molecule XL-784 we used advanced assays measuring coagulation (thromboelastography^34^) and haemorrhage (*in vivo* chicken embryo model^35^), respectively. *E. romani* venom (60 ng) induced rapid clot formation in blood from healthy human donors (clotting time of 3.7 mins [SD 0.7] vs 12.4 mins [SD 0.4], non-venom control) and induced a hypercoagulable state evidenced by increased maximum clot firmness (MCF) (65.7 mm [SD 0.6] vs 58.8 mm [SD 3.0]) (Figure 3c). While XL-784 had no effect on clotting time and only a modest reduction of venom-induced MCF (28.9% reduction), DC-174 prolonged the mean clotting time (venom, 3.7 mins [SD 0.7] vs venom and DC-174, 8.6 mins [SD 0.7]) and returned the MCF to the normal range (mean of 60.3 [SD 0.6] vs 58.8 [SD 3.0] non-venom control, compared to 65.7 mm [SD 0.6] venom only control) (Figure 3c). To assess protection against venom-induced haemorrhage in an *in vivo* vascularised system, groups (n=5) of six-day-old chicken embryos were topically dosed onto the vitelline vein with *E. romani* venom (10 µg/egg), followed by drug (1 µg/egg). Embryos dosed with venom died within 6 hours (Figure 3d) and showed clear signs of haemorrhage as early as 1 hr post-dosing (Figure 3e). While XL-784 only provided partial protection against envenoming, with three (60%) embryos surviving until the 6 h experimental endpoint, treatment with DC-174 resulted in complete survival (Figure 3d) and also prevented venom- induced haemorrhage (Figure 3e).

### Oral dosing of DC-174 shows efficacy against envenoming *in vivo*

We next progressed evaluation of DC-174 into a modified version of the WHO-recommended, *in vivo* neutralisation of venom-induced lethality murine model of snakebite envenoming^14,36^. In these rescue experiments, groups of mice (n=5) received a previously established lethal challenge dose of *E. romani* venom (90 µg^14^) intraperitoneally, followed by a 20 mg/kg oral dose of DC-174 in standard suspension vehicle (SSV). A venom control group received SSV only instead of drug. All mice in the venom-only group succumbed to the lethal effects of envenoming within 161 mins (111-161 mins, mean 138 mins), defined as implementation of euthanasia when humane endpoints associated with severe systemic envenoming were observed. Compound DC-174 significantly prolonged survival times (*p* = 0.01), with three mice succumbing to the lethal effects of the venom much later in the experimental timeframe (272, 370 and 372 minutes), while the remaining two mice survived until the 8-hour experimental endpoint (Figure 4a).

**Figure 4.**
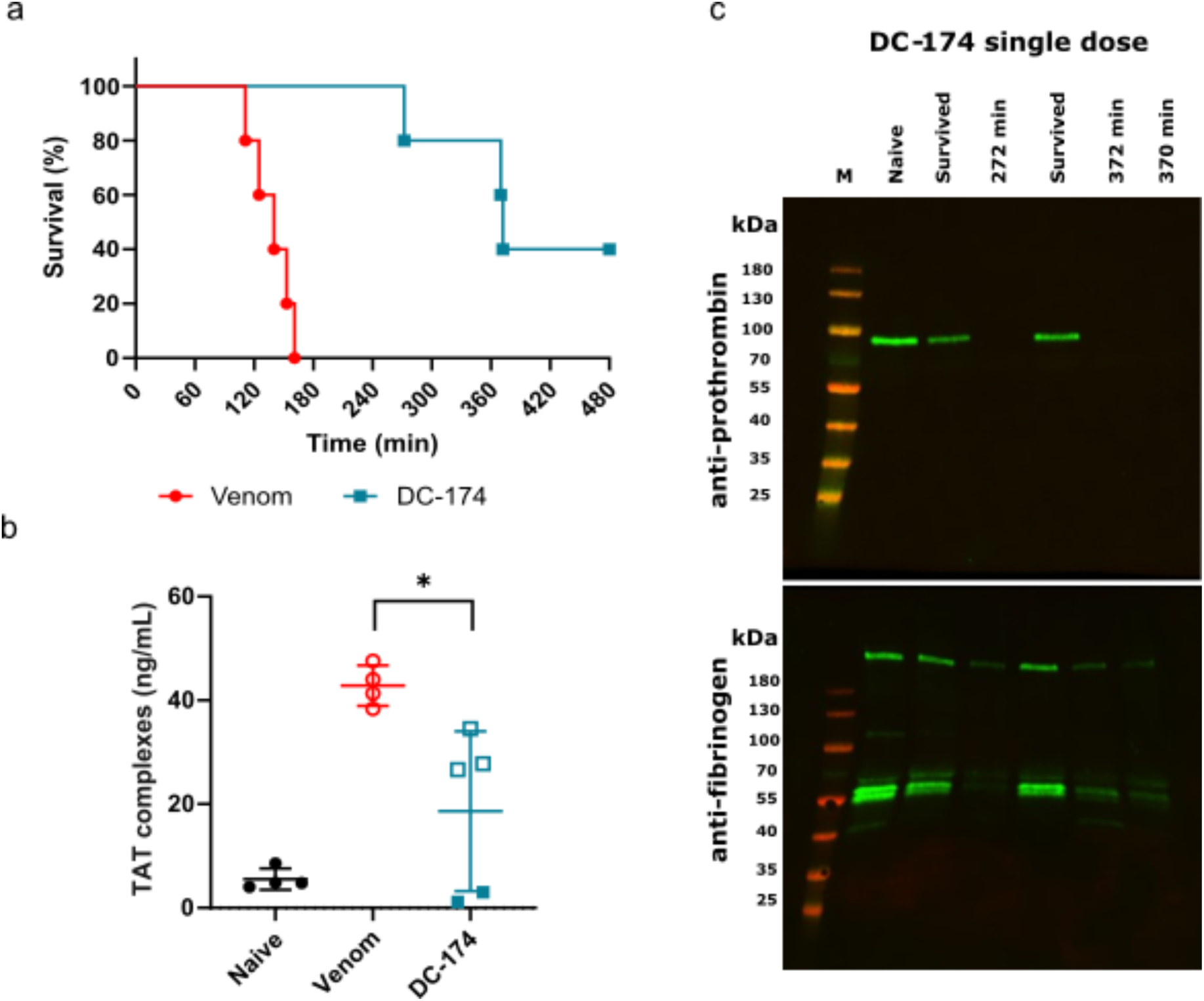
Oral DC-174 prolongs survival in a murine preclinical model of severe envenoming. a) Kaplan-Meier survival graph for mice (n=5) dosed with *E. romani* venom (90 µg intraperitoneally) followed by immediate oral dosing with DC-174 (20 mg/kg, teal squares) or SSV drug vehicle control (red circles). b) ELISA quantification of TAT complexes from murine plasma samples from (a). Data from naïve non-dosed mice (n=4) are shown for comparison, and open and closed squares are used to discriminate between animals dosed with DC-174 that died or survived, respectively. Error bars represent means ±SD. An ordinary one- way ANOVA was performed for all groups with significance indicated by an asterisk (F(2,10) = 14.25, *p =* 0.012). c) Anti-fibrinogen and anti-prothrombin western blots on murine plasma samples from (a) (n=5), compared with samples from a naïve control mouse. Where animals succumbed to venom effects, they are annotated with time to death. M – molecular weight marker.

Blood samples were collected post-euthanasia for all experimental animals (at 8 hours for survivors) and plasma isolated for biomarker analysis. We quantified thrombin-antithrombin (TAT) complexes, which are a marker of thrombin generation (i.e. proxy for coagulopathy) that has been shown to correlate with severity of envenoming^14^. Resulting TAT levels were highly elevated in venom-only controls (mean 42.8 ng/mL vs 5.5 ng/mL in non-envenomed mice). Mice receiving oral DC-174 displayed significant reductions in TAT levels (*p* = 0.01), with those that died during the experiment exhibiting modest reductions compared with the venom-only controls (mean 29.6 ng/mL vs 42.8 ng/mL, respectively), while the two survivors had TAT levels comparable to those of non-envenomed mice (mean 2.1 ng/mL vs 5.5 ng/mL, respectively), suggesting protection against coagulopathic venom effects. Visualisation of murine plasma prothrombin and fibrinogen via Western blotting showed a similar pattern. Venom only control mice had plasma depleted of prothrombin and almost completely degraded fibrinogen (Supplementary figure 4). Mice dosed with oral DC-174 that succumbed to envenoming showed similar profiles to the venom-only controls, with only traces of fibrinogen visible, but those that survived displayed intact prothrombin and fibrinogen profiles, comparable with those of naïve mice (Figure 4c).

### A pharmacokinetically-informed two dose oral regimen improves the *in vivo* efficacy of DC-174

While oral dosing of DC-174 significantly prolonged murine survival, the observed protection against severe envenoming was modest (257 min increase in mean survival times), which prompted us to evaluate the pharmacokinetic (PK) properties of the drug *in vivo*. PK analysis was evaluated over 24 hours post-oral administration (20 mg/kg) of DC-174 in mice (n=9). Resulting data showed DC-174 had a rapid time to maximal drug concentration (T_max_, 0.5 h), but only a moderate half-life (T_1/2_, 1.41 h) and exposure (C_max_, 311 ng/mL) (Figure 5a). The moderate drug exposure could be the result of the low logD_7.4_ of DC-174 (logD_7.4_ of 0.2), which can limit drug permeability and absorption^37,38^. These data may explain the drop off in efficacy observed 4 h after oral dosing, since drug levels are predicted to be 26.1 ng/mL at this timepoint, an amount that equates to only 8.4% of the C_max_.

**Figure 5.**
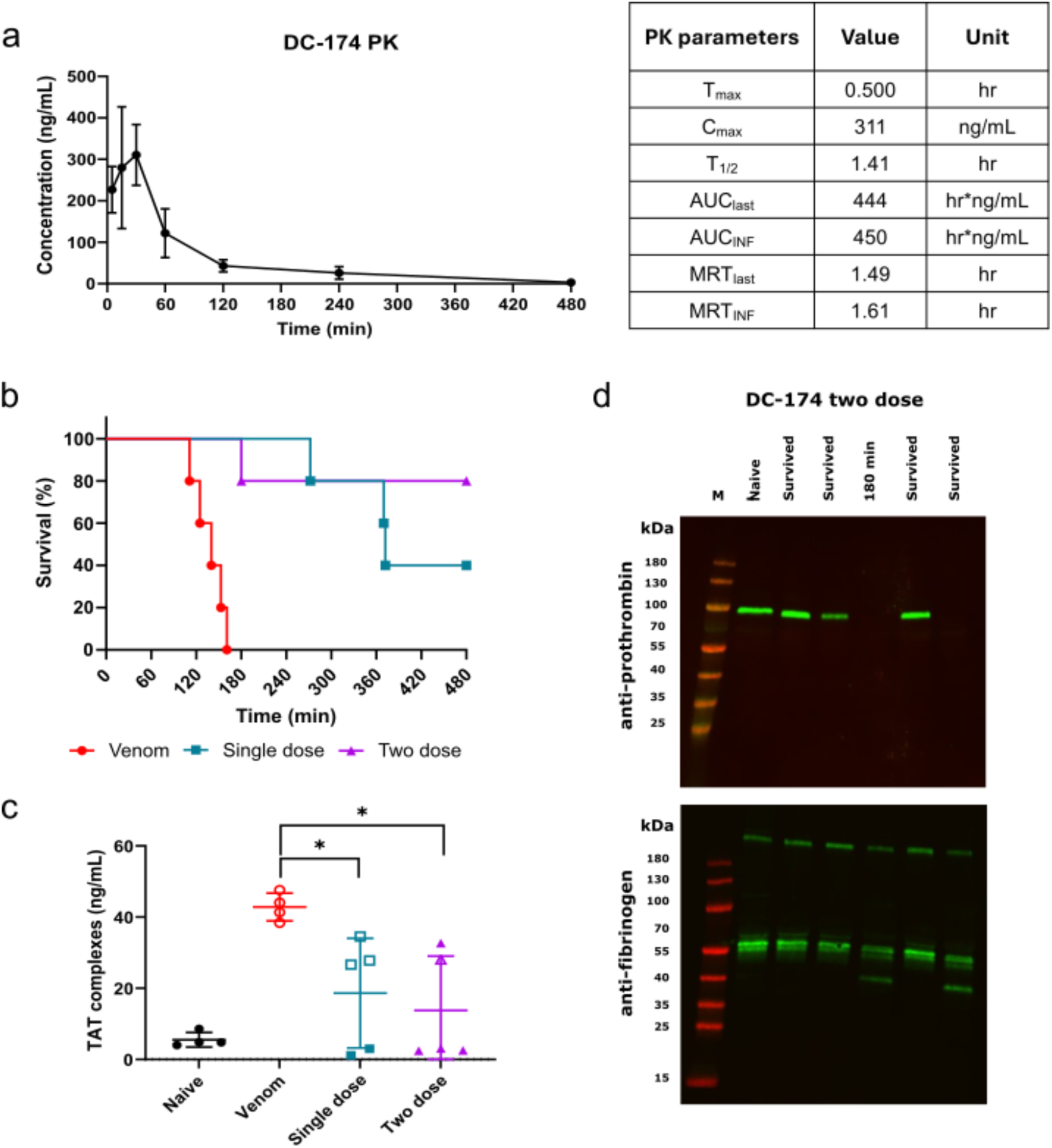
A two dose oral dosing regimen of DC-174 outperforms single oral dosing in a murine preclinical model of severe envenoming. a) PK profiles of mice (n=3) dosed orally with 20 mg/kg DC-174. b) Kaplan-Meier survival graph for mice (n=5) dosed with *E. romani* venom (90 µg intraperitoneally) followed by oral dosing of DC-174 (20 mg/kg) immediately (single dose regimen) or immediately followed by a second 20 mg/kg dose 1.5 h later (two dose regimen). The venom vehicle controls received venom and a single dose of SSV drug vehicle control immediately thereafter. c) ELISA quantification of TAT complexes from murine plasma samples from (b). Data from naïve non-dosed mice (n=4) are shown for comparison, and open and closed squares and triangles are used to discriminate between animals dosed with DC-174 that died or survived, respectively. Error bars represent means ±SD. An ordinary one-way ANOVA was performed for all groups with significance indicated by asterisks (*, F(3,14) = 7.517, *p<*0.05; venom vs single dose, *p* = 0.037, venom vs. two dose, *p* = 0.011). d) Anti-fibrinogen and anti-prothrombin western blots on mouse plasma samples collected from the repeat dosing of DC-174 (n=5) vs a naïve control. Where animals succumbed to venom effects, they are annotated with time to lethality. M – molecular weight marker.

To test this hypothesis, we repeated the *in vivo* murine efficacy study using a multiple oral dosing regimen, with mice (n=5) receiving the same 20 mg/kg oral dose of DC-174 immediately after venom challenge, followed by a second oral 20 mg/kg dose 1.5 h later. This multiple dosing regimen conferred an increase in efficacy, with only one experimental animal succumbing to the lethal effects of the venom (at 180 mins), while the remaining four animals survived the duration of the 8 h experiment (Figure 5b). Mean survival times increased from 138 mins (venom only control) to 420 mins (two dose oral regimen). Analysis of resulting plasma samples revealed a further decrease in TAT levels when DC-174 was dosed twice (mean 13.7 ng/mL), lower than those observed with a single dose (mean 18.6 ng/mL) (Figure 5c). The number of mice with intact plasma prothrombin and fibrinogen also increased following repeat dosing of DC-174 (Figure 5d). Prothrombin was detectable in three of the four surviving mice, while fibrinogen was detectable in all multiply dosed animals, and with noticeably reduced levels of degradation compared with those dosed once (Figure 4c and Figure 5d). Collectively, this data suggests that the half-life of DC-174 may limit its efficacy, and we show that repeated oral administration provides a route to circumvent this limitation, resulting in increased efficacy against snakebite envenoming *in vivo*.

### Predicting DC-174-SVMP binding interactions by molecular modelling

A major challenge in developing small molecule treatments for snakebite is achieving broad- spectrum activity across different toxin isoforms and venoms. Our *in vitro* results show that compound DC-174 meets these criteria. To understand its binding, and that of the parent compound XL-784, molecular docking was performed using X-ray crystal structures of various SVMPs, including P-I and P-III classes (Figure 6 and Supplementary figure 5). No P-II SVMP structures were available for this analysis.

**Figure 6.**
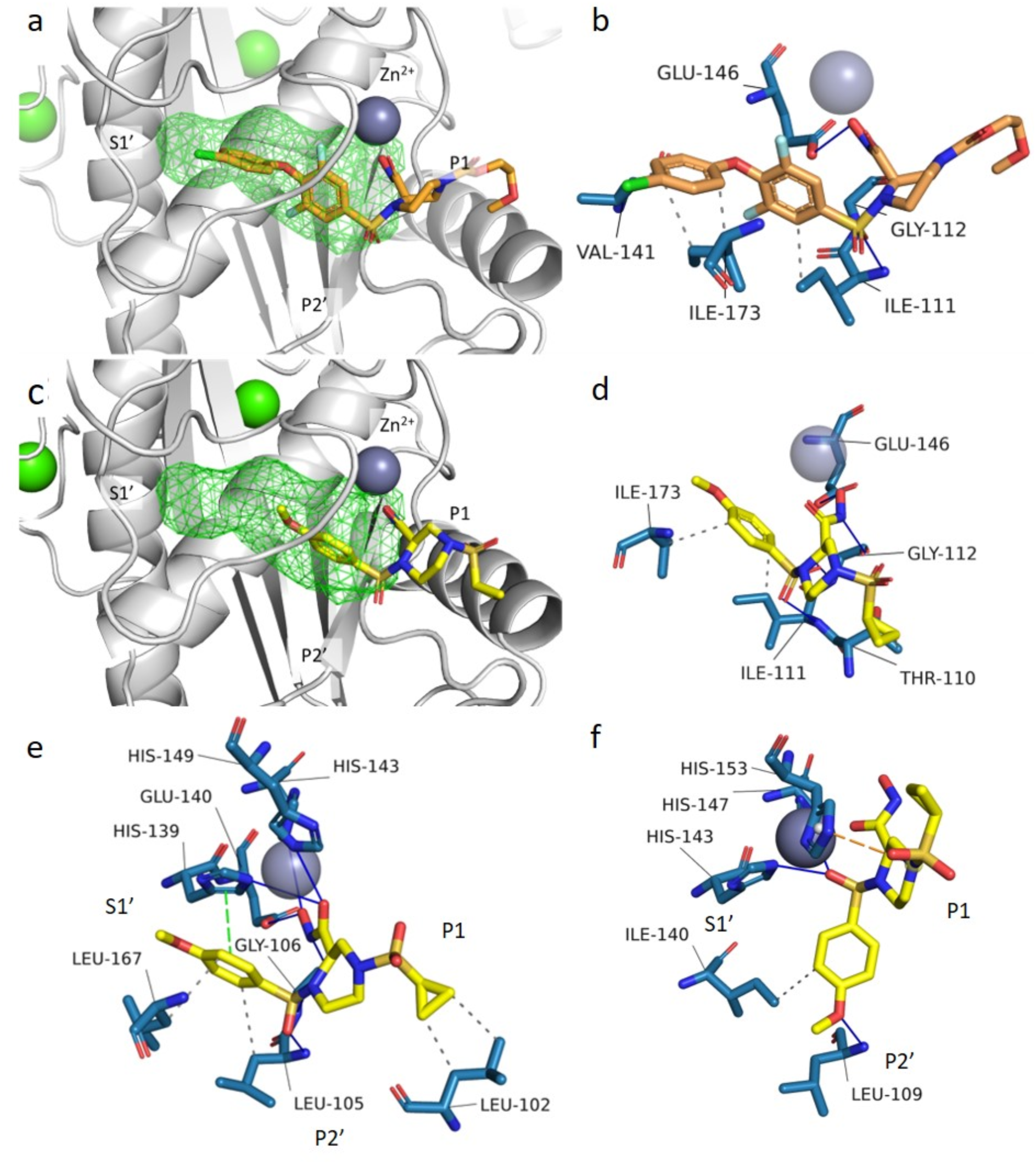
Molecular modelling predicted binding poses and interactions between SVMPs with XL-784 and DC-174. Docked poses of XL-784 (orange sticks in a and b), and DC-174 (yellow sticks in c and d) in the P-III SVMP bothropasin (PDB: 3DSL). e) Docked poses of DC-174 in RVV-X (PDB: 2E3X). f) Docked poses of DC-174 in ecarin (PDB: 9CLP) SVMPs. a) and c) Structure of the docked inhibitors in the SVMP binding site (grey cartoon), with the S1’ pocket highlighted in green wireframe. Zn^2+^ and Ca^2+^ ions are shown in deep blue and green, respectively. b), d), e) and f) Analysis of non-covalent interactions formed by inhibitors in the docked structures. Binding residues are shown in blue, hydrophobic interactions are shown as grey dashed lines, π-cation interactions are shown as orange dashed lines, π-stacking interaction are shown as green dashed lines, and hydrogen bonding interactions are shown as blue solid lines.

We explored structure-activity relationships (SAR) assuming XL-784 binds similarly to prinomastat. Molecular docking of XL-784 in the P-III SVMP bothropasin supported this (Figure 6a and 6b), showing a binding pose where its hydroxamic acid coordinates with the catalytic Zn²⁺ and its hydrophobic biaryl ether fits into the S1’ pocket^39^, consistent with prinomastat binding to MMP13. The binding pocket primarily involves hydrophobic interactions with aliphatic residues Ile111, Val414, and Ile173. Hydrogen bonds form between the sulfone oxygen and hydroxamic acid with nearby backbone NH, carbonyl groups, and a glutamate in the Zn²⁺ binding domain. In the P-I SVMP atrolysin C, similar interactions occur, along with halogen bonding to backbone carbonyls and π-stacking with His142 in the Zn²⁺ domain (Supplementary figure 5a and 5b). In the P-I SVMP BaP1, although the core piperazine H-bonding is conserved, the S1’ pocket is too small for the biaryl ether of XL-784, which instead extends into the P2’ pocket (Supplementary figure 5c and 5d). This behaviour is also seen in the P-III SVMP ecarin, where a stabilizing salt bridge forms (Supplementary figure 5e). In the P-III SVMP RVV-X, XL-784 adopts a reversed pose, avoiding the S1’ pocket (Supplementary figure 5f). These findings across SVMP isoforms suggest that the size of the S1’ substituent strongly influences the binding mode and inhibitory spectrum of the molecule.

Compound DC-174 was docked into SVMP crystal structures to compare its interaction profile with XL-784. In bothropasin (Figure 6c and 6d), the hydrogen bonding between the piperazine- hydroxamic acid and key residues, including glutamate in the Zn²⁺ binding domain, was preserved with additional interactions with sulfone and backbone NH groups observed. The methoxyphenyl ring of DC-174 fits well into the S1 pocket, forming hydrophobic interactions with Ile173 and Ile111, while all key hydrogen bonds from XL-784 were conserved with additional ones formed. Docking studies of DC-174 in BaP1 (Supplementary figure 6a and 6b), atrolysin C (Supplementary figure 6c and 6d), and RVV-X (Figure 6e) show that its truncated monoaryl ring is well tolerated across multiple SVMP isoforms, unlike XL-784, which was hindered by the small S1’ pocket in BaP1 (Supplementary figure 5c). In ecarin, DC-174 adopted an outward pose along the P1 and P2’ pockets, stabilized by π-cation and hydrogen bond interactions with Leu109 (Figure 6f). However, its piperazine-hydroxamic acid hydrogen bonding was not conserved, instead forming a bond via the sulfone group. Interaction analysis indicates that key hydrogen bonding motifs are conserved in both P-I SVMP isoforms (Supplementary figure 6b and 6d). Unique π-stacking interactions with His142 was only observed in BaP1 (Supplementary figure 6b). Unlike XL-784, the cyclopropyl sulfonamide moiety in DC-174 forms hydrophobic interactions with residues in atrolysin C, BaP1, and RVV-X, and an additional hydrogen bond with His152 in atrolysin C.

Further docking and molecular dynamic studies (see Supplementary information for molecular modelling; Supplementary figure 7 and 8; Supplementary table 4 and 5) collectively provide a structural basis for the broader inter-species and cross-isoform inhibitory activity of DC-174 over its parent compound XL-784. The predicted binding mode of DC-174 across structurally diverse SVMPs also indicates its venom-neutralizing effect stems from direct SVMP inhibition.

## Discussion

Herein, we have described the first experimental design, synthesis and evaluation of SVMP inhibitory properties, resulting in an orally bioavailable lead candidate snakebite drug. We showed that a rationally designed small molecule inhibitor can effectively neutralise diverse SVMP toxins from a wide range of medically important snake venoms. Unlike the parent molecule XL-784 and other previously described SVMP inhibitors (i.e. prinomastat)^24,40,41^, inhibitor DC-174 neutralises all three of the structurally diverse SVMP sub-classes (i.e. P-I, P- II and P-III), likely as the result of its truncated monoaryl ring system. Further, inhibitor DC- 174 is capable of fully or near fully preventing the generation of thrombin by inhibiting prothrombin proteolysis by *E. romani* P-I and P-III SVMPs, respectively (Figure 3). The ability of certain SVMPs to generate thrombin and activate the clotting cascade forms the basis of venom-induced consumption coagulopathy (VICC), whereby venom metalloproteinases cleave and deplete the internal pool of clotting factors leading to incoagulable blood. As such, DC-174 can prevent the initiation of one of the main pathways responsible for causing coagulopathy, which in turn exacerbates haemorrhage, in snakebite patients. These findings are reinforced from the *in vivo* models applied, with DC-174 preventing haemorrhage in an *in vivo* egg embryo model and significantly prolonging survival and reducing coagulopathy when administered orally following *E. romani* venom challenge in a murine model of envenoming (Figures 3-5).

The murine PK data obtained for DC-174 shows that it only takes 30 minutes for drug levels to reach their peak following oral dosing, but that the half-life is rather short at 1.41 h (Figure 5a). The clearance of DC-174 may underlie the intermediate level of protection observed following single oral administration in mice (i.e. significant increase in survival times, but only 40% survival at 8 h). In agreement with this hypothesis, we showed that repeated administration of DC-174, with a second dose 1.5 h after the first, improves drug efficacy, resulting in 80% survival at the end of the 8 h experiment. Plasma biomarkers of coagulation also demonstrated that repeat dosing of DC-174 increased efficacy, with TAT, prothrombin and fibrinogen levels superior to those observed with single dosing.

To contextualise these findings, the metal chelator DMPS - an SVMP inhibiting molecule that is currently in clinical trials for snakebite indication^42^ - is the only other SVMP inhibiting drug with demonstrated efficacy via oral dosing^14^. Here we show that DC-174 outperforms DMPS both in terms of *in vitro* potency (ERO EC_50_s, 9.7 nM vs 1.65 µM)^14^ and *in vivo* efficacy (oral dosing murine survival of 80% vs 40% at 8 h with 20 mg/kg twice dosing vs 30 mg/kg single dosing, respectively^14^). Collectively these results underscore the increased efficacy of DC-174 relative to its parent molecule, XL-784 and other lead SVMP-inhibiting drugs, and emphasises the importance of applying rational, structure-informed chemical modification to improve PK and PD parameters of next-generation treatments for snakebite envenoming.

This work establishes a novel drug discovery pipeline for SVMP inhibitors that provide reassurance of “on-target” mechanism of action. Further, the highly efficient chemical synthetic strategy for production of DC-174 only required three chemical reaction steps and utilised affordable, commercially available materials for synthesis, highlighting the potential future cost-effectiveness of oral drugs as alternatives to the costly polyclonal antibody-based antivenom therapies currently available. Additional development work on DC-174 will next be required to establish robust preclinical toxicology profiles and expand PK profiling to predict a suitable human dosing regimen. In parallel, our future medicinal chemistry efforts will focus on maximising the metabolic stability characteristics of this scaffold to provide a broad-spectrum efficacious molecule compatible with a single dose regimen, as well as evaluating the potential of DC-174 as a component of combination therapies with other oral drugs that target different venom toxins^15,16^, such as the PLA_2_ inhibitor varespladib^19^.

DC-174 broadens the current narrow portfolio of only two SVMP inhibiting drugs progressing into clinical development for snakebite, thus helping to avoid attrition as the field moves towards an optimised oral treatment. This novel SVMP inhibitor represents one of several unique molecules currently being developed through our discovery platform, with the chance of additional candidates entering full development for clinical progression within the next two years, with the long term goal of accelerating treatment options available for the tropical communities most afflicted by life-threatening snakebite.

## Methods

### Chemistry

Chemicals were purchased from Sigma-Aldrich, Fluorochem, or Alfa Aesar and used without further purification. Unless otherwise stated, anhydrous solvents and dry atmospheric conditions were used in all reactions. Unless otherwise stated, all reactions were carried out at room temperature (rt) under atmospheric pressure. Thin layer chromatography (TLC) was carried out on silica gel plates 60 F_254_ and visualized under UV light at 254 nm. Potassium permanganate dip was used to visualize non-UV active compounds. Flash column chromatography separations using 40−63μm silica were performed by the gradient elution method, and the elution solvent system is given in each instance. Reverse-phase high-performance liquid chromatography (RP-HPLC) was carried out on an Agilent 1260 Infinity system, using a ZORBAX SB-C18 (9.4 mm x 250 mm, 5 µm) column at a rate of 4 mL/min for semipreparative RP-HPLC and a ZORBAX Eclipse Plus C18 (4.6 mm x 100 mm, 3.5 µm)) column at a rate of 1 mL/min for analytical RP-HPLC. The following mobile phases were used A, H_2_O (+ 0.1% v/v formic acid), and B, ACN (+ 0.1% v/v formic acid). The standard analytical method for analytical RP-HPLC (for which retention times of compounds are given) was 5% B for 1 min and a linear gradient from 5% to 95% A from 1 to 12 min (followed by a 3 min hold at 95% B and then a 1 min linear gradient from 95% to 5% B and a 1 min re-equilibration at 5% B). All final compounds were >95% RP-HPLC analytically pure as assessed by peak integration at 254 nm. A typical semipreparative HPLC method was carried out as described: 5% B for 1 min and a linear gradient from 5% to 95% B from 1 to 12 min (followed by a 2 min hold at 95% B and then a 1 min linear gradient from 95% to 5%). High-resolution electrospray ionization mass spectra (HRMS-ESI) were recorded on an Agilent QTOF 6540 spectrometer. Proton (^1^H) and carbon (^13^C) nuclear magnetic resonance (NMR) spectroscopy were carried out using either a Bruker AMX 400 (^1^H, 400 MHz; ^13^C, 101 MHz) or 500 MHz (^1^H, 500 MHz; ^13^C, 126 MHz) spectrometer. Chemical shifts are listed on the δ scale in parts per million, referenced to CDCl_3_ (^1^H NMR δ7.26; ^13^C NMR δ77.16) or MeOD-*d_4_* (^1^H NMR δ3.31; ^13^C NMR δ49.03) with residual solvent as the internal standard and coupling constants (*J*) recorded in hertz. Note - that not all magnetically non-equivalent carbons were observed in the ^13^C NMR spectrum for all compounds. Signal multiplicities are assigned as follows: s, singlet; d, doublet; t, triplet; q, quartet; dd, doublet of doublets; br, broad; m, multiplet. Single-crystal X-ray diffraction was mounted on a MiTGen tip via Parbol oil and data were collected on a Bruker D8 Venture Photon III diffractometer. The crystal was kept at 200.0 K during data collection. Using Olex2^43^, the structure was solved with the XT structure solution program using Intrinsic Phasing and refined with the XL refinement package using Least Squares minimisation.

Crystallographic data for DC-174 have been deposited in CCDC database (Deposition number: 2451930).

### Molecular Modelling and Docking

To investigate binding of XL-784 and DC-174 in representative SVMPs, we carried out molecular docking simulations. Compounds were built and energy minimised at the MMFF level using Spartan24 [Spartan ’24 version 1.1.0 (Wavefunction, 2024)]. Energy minimised compounds were docked into available crystal structures of relevant SVMP from venoms used in *in vitro* profiling; *Bothrops jararaca* (bothropasin, PDB: 3DSL)^44^, *Crotalus atrox* (atrolysin C, PDB: 1ATL)^45^, *Bothrops asper* (BaP-1, PDB: 2W13)^46^, *Echis carinatus* (ecarin, PDB: 9CLP)^47^ and *Daboia siamensis* (RVV-X, PDB: 2E3X)^48^. Docking was carried out in GOLD^49^ with a docking protocol validated by re-docking of the co-crystallised ligands for each SVMP. Docking was carried out without allowed early termination and allowed flipping of amides, pyramidal nitrogen and flipping ring corners, default settings were used otherwise. Harmonic distance restraints were applied to atoms involved in chelation of the catalytic zinc in the SVMP catalytic site with minimum distance set at 1.5 Å, maximum distance set at 2.5 Å and spring constant set at 10 – in each extracted ligand the bound moieties are carboxylate groups interacting with zinc by each oxygen. The CHEMPLP scoring function to generate poses. This afforded docking poses with RMSD values of < 1.5 Å in each structure with respect to the extracted ligand.

XL-784 and DC-174 were docked into each SVMP using the validated protocol. Harmonic restraints were applied between the hydroxamic acid carbonyl and sp^3^ oxygen, and Zn with the same distance thresholds and spring constant as described above. Top ranked poses by CHEMPLP score in each SVMP were extracted and analysed for non-covalent interactions using PLIP^50^. Further computational details are provided in the supporting information.

### Biology

Venoms were extracted from specimens of *E. romani* (Nigeria [formerly *E. ocellatus*]), *C. atrox* (United States) and *B. arietans* (Nigeria) maintained within the herpetarium at the Centre for Snakebite Research and Interventions (CSRI) at the Liverpool School of Tropical Medicine (LSTM). Crude venoms were pooled by species and lyophilised for long term storage at 2-8°C. Venoms from *C. rhodostoma* (Thailand) and *B. jararaca* (Brazil) were sourced from the historical collection of lyophilised venoms held at LSTM. Venoms were reconstituted to 10 mg/mL in sterile phosphate buffered saline (PBS, pH 7.4) (cat. no. 10010-015, Gibco) prior to use.

### *In vitro* inhibition of enzymatic SVMP activity

Compounds of interest were stamped onto 384-well plates at a volume of 0.91 µL at a concentration of 1 mM in 100 % DMSO (D2650, Sigma Aldrich). Positive and negative control wells were stamped with 0.91 µL of 100 % DMSO. To the test wells and the venom-only positive control wells, 15 µL of venom diluted to 0.066 µg/ml in PBS was added, giving a final amount of 1 µg/well. In the no venom negative control wells, 15 µL of PBS was added. The plates were then sealed, span briefly in a Platefuge (C2000, Benchmark Scientific) and incubated at 37 °C for 25 minutes to allow venom-inhibitor interaction. The plates were removed from the incubator for 5 minutes to allow acclimation to room temperature, then 75 µL of Mca-KPLGL-Dpa-AR-NH2 fluorogenic peptide substrate (ES010, BioTechne), diluted to a concentration of 9.1 µM in SVMP substrate buffer (150 mM NaCl, 50 mM Tris HCl pH 7.5), was added to every well using a ViaFlo 384 (Integra). The final assay volume was 91 µL, yielding a final drug concentration of 10 µM, a final DMSO concentration of 1 %, and a final SVMP substrate concentration of 7.5 µM. The plates were then read kinetically using a fluorescence intensity protocol on a CLARIOStar microplate reader (BMG Labtech) at 320 nm excitation, 420 nm emission, for 60 minutes. At a timepoint appropriate for the venom to have cleaved all substrate in the well (*E. romani* and *C. rhodostoma* 20 min; *B. jararaca* and *C. atrox* 30 min; *B. arietans* 40 min), raw fluorescence values were normalised to be a percentage of the positive and negative control values, with complete inhibition of SVMP activity displaying as 100 % inhibition and no inhibition of SVMP activity displaying as 0 % inhibition. Dose response plates were conducted identically and the % inhibition values at each dose were used to draw EC_50_ curves to estimate the EC_50_ value of each compound tested. All SVMP experiments were repeated independently a minimum of three times.

### *In vitro* inhibition of coagulopathic venom activity

This assay consists of pre-incubating venom (or venom vehicle control) at a species-specific dose with the compound of interest (or compound vehicle control), then adding citrated bovine plasma to the wells and activating the clotting cascade with calcium chloride. The selected venoms (*E. romani* and *B. jararaca*) are procoagulant, resulting in clot formation within 10 minutes compared to ∼30-45 minutes with recalcified normal plasma. As *E. romani* is a more potently procoagulant venom, only 10 ng per reaction was required to initiate rapid clotting, whereas 100 ng of *B. jararaca* venom was required to induce a similarly rapid clotting curve. Compounds of interest were stamped onto 384-well plates at a volume of 0.5 µL and a concentration of 10 mM in 100 % DMSO. Positive and negative control wells were stamped with 0.5 µL of 100 % DMSO. To the test wells and the venom-only positive control wells, 10 µL of venom diluted to 1 µg/ml (*E. romani*) or 10 µg/ml (*B. jararaca*) in PBS was added, giving a final well amount of 10 ng (*E. romani*) or 100 ng (*B. jararaca*) per well. In the no venom negative control wells, 10 µL of PBS was added. The plates were then sealed, span briefly in a Platefuge and incubated at 37 °C for 25 minutes to allow venom-inhibitor interaction. While the plates were incubating, sufficient citrated bovine plasma (S0260, BioWest) to allow 20 µL/well was centrifuged at 3120 RCF for 5 minutes to remove any residual cellular debris and the supernatant was collected. The 384-well plates were removed from the incubator for 5 minutes to allow acclimation to room temperature, then 20 µL/well of 20 mM CaCl_2_ was added and immediately followed by the addition of 20 µL/well of centrifuged citrated bovine plasma, both via a Multidrop Combi reagent dispenser (5840330, ThermoFisher Scientific). The plates were then read kinetically using an absorbance protocol on a CLARIOStar microplate reader at 595 nm for 60 minutes. The time, in minutes, required to achieve half the maximal absorbance value was calculated for each well using Prism (Graphpad, v10.2.3), and then were normalised to be a % of the positive and negative control values, with complete inhibition of procoagulant activity displaying as 100 % inhibition and no inhibition of procoagulant activity displaying as 0 % inhibition. Dose response plates were conducted identically and the % inhibition values at each dose were used to draw EC_50_ curves to estimate the EC_50_ value of each compound tested. All coagulation experiments were repeated independently a minimum of three times.

### Degradation of coagulation factors

Evaluation of SVMP toxin activity of human prothrombin and fibrinogen and inhibition by test compounds was performed by SDS-PAGE degradation gel electrophoresis using venom-purified toxins. P-I, P-II and P-III SVMPs were isolated using an adaptation of the method of Howes et al.^51^. Whole *E. romani* venom (15 mg) was initially separated using size exclusion chromatography (SEC) on a 120 mL column of Superdex 75 (Cytiva). The P-II SVMP (22 kDa) was pure and ready to use at this stage. P-I and P-III SVMP containing peaks were separately applied to a 1 mL high resolution hydrophobic-interaction chromatography (HIC) column (SOURCE 15PHE, Cytiva). The proteins were separated using a 10-column volume gradient of 1.0 M ammonium sulphate in 50 mM Tris-Cl, pH 8.8 to 25% ethylene glycol in 50 mM Tris-Cl, pH 8.8. The P-I eluted as two peaks of proteins with identical molecular weight (21 kDa) which were combined back together. The P-III-containing fraction from SEC eluted from HIC as multiple peaks of P-III SVMP isoforms, well separated from the earlier eluting L-amino acid oxidase and high molecular weight C-type lectin-like proteins. The proteins in the PIII-containing peaks were pooled back together. Finally, both the P-I and P-III SVMPs were dialysed into TBS (25 mM Tris-Cl, pH 7.8, 0.15 M NaCl) ready for degradation assays.

Prothrombin degradation was measured by incubating 2 µg of human prothrombin (Haematologic Technologies INC) in the presence or absence of drug and the P-I, P-II, and P- III SVMPs. As the P-II SVMP did not cleave the prothrombin substrate, it was excluded from inhibitory assaying. Briefly, drug or DMSO and 200 ng of purified SVMP isoform were incubated for 30 min at 37 °C in a water bath, after which 2 µg of human prothrombin were added and the samples incubated for another hour at 37 °C. Drug concentrations used were 2 µM for P-I and P-III SVMPs. Samples were then processed for loading onto a 4-20% SDS-PAGE gel (Biorad) and were run under reducing conditions for 45 min at 180 V. The gels were stained with Coomassie Brilliant Blue R250 and degradation profiles were visualised and images captured using a GelDoc (Biorad).

Degradation of human fibrinogen was measured in a similar manner. Human fibrinogen (Merck) was added at 5 µg/reaction and incubated with the different SVMP isoforms in the presence or absence of drugs. Drug concentrations used were 10 µM for P-I and P-II SVMP and 400 nM for P-III SVMP. The reactions consisted of 200 ng SVMP isoform and the appropriate concentration of drug, which were then incubated for 30 min at 37 °C in a water bath. Five micrograms of fibrinogen were then added and the samples incubated for another hour at 37 °C, before separation by SDS-PAGE as described above.

### Thromboelastography

We measured the coagulation profile of whole human blood using thromboelastography, and compared with venom stimulated, and venom and drug stimulated, clotting profiles. Blood samples were obtained according to ethically-approved protocols (LSTM research tissue bank, REC ref. 11/H1002/9) from consenting healthy volunteers who confirmed they had not taken any anticoagulant treatments for at least three months prior to blood collection. Blood samples were collected in ACD-A blood tubes. Using a ROTEM delta (Werfen) a final volume of 300 μL blood per reaction was pipetted into prewarmed cups (37 °C), with four channels per run. Whole blood was warmed for 5 mins at 37 °C, during which time 12 μL of *E. romani* venom (5 ng/nL concentration, 60 ng dose), 15 μL of DMSO vehicle or test compounds (5 μM final concentration, 0.2 % DMSO), and 20 μL of Startem reagent (CaCl2, Werfen) was prepared. The maximum incubation time of venom and inhibitors was 2 minutes, before 253 μL of whole blood were added and mixed before reading on the ROTEM delta. The commonly used parameters of clotting time and maximum clot firmness (MCF) were reported, from a minimum of three independent measurements per experimental group.

### Chicken egg embryo model

To compare the *in vivo* efficacy of DC-174 with XL-784, a chicken embryo model was applied^35^. Fertile hens eggs were supplied on day 1 post fertilisation from UK supplier MedEggs Ltd. Eggs were immediately placed horizontally to incubate at 37 °C until day 5 of development. Candling was conducted to mark the position of the embryo to guide windowing. Any infertile eggs were removed at this stage. To access the embryos, all fertile eggs were then windowed under laminar flow. In brief, 6 – 8 mL of albumin was removed using a 23G needle. The area marking the position of the embryo was then covered with medical tape, and a dish of shell removed using sharp dissection scissors. The exposed embryo was then covered using parafilm and the egg returned to the incubator overnight. Surviving embryos were then randomly allocated into groups of n = 5. Embryos then received either *E. romani* venom 10 µg/egg or *E. romani* venom immediately followed by XL-784 or DC-174 (1 µg/egg). Treatments were given in a total volume of 10 µL, administered directly onto (i.e. topically) the vitelline vein. Embryos were returned to the incubator immediately following dosing, and then observed at 1, 2, 4 and 6 hours. Survival was monitored via the observation of a heartbeat. Venom pathology was captured using a Motic SMZ 171-TLED microscope system. In brief, at each observation time point (1, 2, 4 and 6 hours post envenoming) images were captured using a Moticam X5 Wi-Fi camera. To track changes across the experimental time course, the imaging position was kept consistent at each timepoint for individual embryos.

### *In vivo* protection against venom-induced lethality

Murine preclinical studies were conducted under protocols approved by Animal Welfare and Ethical Review Boards of the Liverpool School of Tropical Medicine and University of Liverpool under project licence (PPL No. P5846F90) approved by the UK Home Office in accordance with the UK (Scientific Procedures) Act 1986. Male CD-1 mice (18-20 g) were purchased from Janvier (France) and acclimatised for 7 days prior to experimentation. Mice were housed in groups of 5 in Tecniplast GM500 cages within a specific pathogen-free facility. Room conditions were set at 20-24 °C and 45 – 65% humidity, 12/12 hour light cycles. Mice were allowed ad lib access to Picolab 5R53 food (Lab Diet, USA) and reverse osmosis water and housed with nesting material and environmental enrichment materials.

Mice weighed 22-28 g at the start of study. 10 mg/kg morphine was administered subcutaneously as an analgesic 15 minutes prior to venom administration. To determine the efficacy of DC-174, a modified version of the WHO neutralisation assay was used^52^. Following analgesia, mice received an intraperitoneal injection of 90 µg *E. romani* venom in 100 µL PBS, corresponding to 5 x the intravenous LD50 dose^53^. DC-174 was solubilised in SSV (carboxymethyl cellulose (0.5%), benzyl alcohol (0.5%), Tween 80 (0.4% v/v) and NaCl (0.9%)) and gently sonicated at 24 °C for 5 minutes. Mice received an oral dose of either 20 mg/kg DC-174 in SSV, or SSV-only (n=5/group, 50 µL/dose), immediately following venom administration. A third group received venom followed by two oral doses of DC-174, one immediately after venom administration and a second identical dose 1.5 h later. No randomisation was used to allocate experimental groups – mice were randomly allocated into cages of five prior to the experiment, and each cage formed one treatment group. No criteria for including or excluding animals was applied, and all data points were included in analyses. Animals were monitored for humane endpoints (loss of righting reflex, seizure, external haemorrhage) for 8 hours to encompass the drug exposure timeframe, and any animals showing such signs were immediately euthanised by rising concentrations of carbon dioxide. All observations were performed by mixed gender experimenters who were blinded to the drug group allocation. Time of death, number of deaths and number of survivors were recorded, where deaths and times of death represent implementation of humane endpoint-dictated euthanasia. Kaplan-Meier survival plots were generated using GraphPad Prism 9.0 (GraphPad Software, San Diego, USA).

### Envenoming biomarker analysis

Blood samples were collected from mice via cardiac puncture immediately following euthanasia. Samples were centrifuged at 1400 xg for 10 mins at 4 °C, and the plasma supernatant stored at –80 °C until use. Plasma TAT levels were determined using a sandwich ELISA provided as a commercial kit (Abcam, ab137994). Mouse plasma samples were diluted 1:150 using Diluent M and were run in duplicate alongside a standard curve. Each well received 50 µL diluted plasma and was incubated for 2 h at room temperature to allow for the reaction with the TAT complexes antibodies coated onto the plate. After five washes, 50 µL of biotinylated mouse TAT complexes antibodies were added to the plate and the reaction allowed to proceed for another 2h. Following another five washes, 50 µL of a streptavidin-peroxidase conjugate were added to the plate and incubated for another 30 minutes. Next, 50 µL of a chromogenic substrate (3,3’,5,5’-tetramethylbenzidine) was added to the plate and the peroxidase-dependent reaction allowed to proceed until a clear blue colour formed in the well. At this point 50 µL of stop solution was added and the absorbance read immediately at 450 nm and 570 nm. The 570 nm absorbance readings were subtracted from the 450 nm ones, and the levels of TAT complexes (ng/mL) extrapolated based on the standard curve. Data was further plotted as means with SD for each mouse sample and n=4-5 samples were used for each experimental condition.

Anti-fibrinogen and anti-prothrombin western blots were performed using mouse plasma to detect the integrity of mouse fibrinogen or prothrombin following envenoming in the presence or absence of DC-174. Mouse plasma samples were diluted 1:50 (anti-fibrinogen blots) and 1:30 (anti-prothrombin blots) and 5 µL was run on 4-20% SDS-PAGE gels (BioRad) at 180V for 1 h. The separated proteins were then transferred onto nitrocellulose membranes using semi-dry transfer on a Trans-Blot Turbo (BioRad) using the following conditions: 1.3 A, 25 V, 7 min. Blots were then stained for 5 min with Revert 700 Total Protein Stain (LICORbio) following the manufacturer’s protocol and imaged on the 700 nm channel on an Odyssey Fc Imaging System to detect protein abundance and ensure equal loading. Membranes were then destained in 0.1 M NaOH, 30% methanol for no longer than 10 min, then blocked in 5% milk TBST for 1 h. The primary antibodies: anti-fibrinogen (ab34269, rabbit polyclonal, Abcam) and anti-prothrombin (PA5-77976, rabbit polyclonal, Thermo Fisher) were diluted 1:1,500 and 1:1,000, respectively in 5% milk TBST. The membranes were incubated with the appropriate primary antibody for 1.5 hours at room temperature, followed by three 5-min washes in TBST. Blots were then incubated for 1 h at room temperature with an IRD 800CW donkey-anti rabbit IgG secondary antibody (1:15000, LICORbio), followed by three 5-min washes in TBS. The blots were imaged on an Odyssey Fc Imaging System at both 700 and 800 nm.

### Drug metabolism pharmacokinetic studies (DMPK) properties assays

The DMPK properties data described in the manuscript were measured via a high-throughput platform by AstraZeneca (United Kingdom). The methods used for the five assays, including logD_7.4_, solubility, plasma protein binding, microsomal and hepatocyte clearance measurements, have previously been reported^54^.

### *In vivo* pharmacokinetic (PK) studies in mice

The *in vivo* PK study was performed by ChemPartner (Shanghai, China). The study protocol is summarized as followed: The PK of compound DC-174 was studied in CD1 male mice (28-29 g, 6-8 weeks) purchased from JiHui Laboratory Animal Co. Ltd (Qualification No.: SCXK (Hu) 2022-0009 20220009018616). Mice had access to food and water throughout the pre- and post-dose sampling period. Compound DC-174 was formulated in 0.5% MC (Sigma, SLCL7954) and 0.1% Tween 20 (Sigma, BCCF2036) in water (RephiLe/S21RDB0304) and administered at 20 mg/kg via oral gavage (N=9). Approximately 110 µL whole blood/time point (sampling at 0.083, 0.25, 0.5, 1, 2, 4, 8 and 24 hours) was collected via serial bleeding in K_2_EDTA tube via facial vein. The blood sample was kept on ice and centrifuged (2,000 xg, 4 °C, 5 min) within 15 minutes post-sampling. The plasma drug concentrations were determined by LC-MS/MS (ESI, Triple Quad 5500 System) on positive ion mode and using dexamethasone as internal standard. Liquid chromatography (LC) was performed using a 2.5 µm Xbridge BEH-C18 column (2.1 mm x 50 mm,) at 60 °C with a rate of 0.6 ml/min. The following mobile phases were used; A: Water (+0.025% Formic acid and 1.0 mM ammonium acetate), B: Methanol (+0.025% Formic acid and 1.0 mM ammonium acetate). The LC method used was 5% B for 0.20 minutes, linear gradient from 5% to 60% B from 0.20 to 1.40 minutes, linear gradient from 60% to 65% B from 1.40 to 2.00 minutes, then hold 90% B from 2.01 to 2.30 minutes, followed by re-equilibration at 5% from 2.31 to 2.60 minutes.

### Statistical information

For EC_50_ data, experiments were performed as independent (biological) repeats at least in triplicate with two technical repeats per biological repeat. EC_50_ curves were generated in Graphpad Prism on data normalised to the positive and negative controls (100% inhibition positive control / 0% inhibition negative control) using the linear regression function. Ranges were calculated as mean ±SEM. Ordinary one-way ANOVAs with Tukey’s multiple comparisons test were performed for ROTEM thrombolelastography data and TAT ELISA data, with statistical outputs reported in the relevant figure legends. All error bars represent the mean ± SD.

## Ethics Statement

Human blood samples were obtained according to ethically-approved protocols (LSTM Research Tissue Bank, LSTM Research Ethics Committee ref. 11/H1002/9) from consenting healthy volunteers who confirmed they had not taken any anticoagulant treatments for at least three months prior to blood collection. Murine preclinical studies of drug efficacy were conducted under protocols approved by Animal Welfare and Ethical Review Boards of the Liverpool School of Tropical Medicine and University of Liverpool under project licence (PPL No. P5846F90) approved by the UK Home Office in accordance with the UK (Scientific Procedures) Act 1986. *In vivo* PK studies conformed to AAALAC International and NIH [Assurance ID: F16-00265 (A5953-01)] guidelines as reported in the Guide for the Care and Use of Laboratory Animals, National Research Council (2011); People’s Republic of China, Ministry of Science & Technology, ‘Regulations for the Administration of Affairs Concerning Experimental Animals’, 1988.

## Author contributions

N.R.C., J.K., P.O.N., N.G.B. conceptualised, acquired funding and administered the project.

D.J.W.C. completed the chemistry, design of compounds, synthetic routes and analysis. A.P.W., L-O.A., and R.H.C. performed *in vitro* bioassays. M.C.W. isolated SVMP toxins. C.D. performed the egg embryo assay. A.E.M., E.C., E.S. and N.R.C. performed the *in vivo* efficacy preclinical studies. L-O.A. carried out envenoming biomarker analysis. C.M.W., N.G.B. carried out molecular modelling. D.J.W.C., L-O.A., A.P.W., R.H.C., C.M.W., R.G., N.M., S.C.L., N.G.B., N.R.C. and P.M.O. wrote the manuscript. All authors edited the manuscript.

## Acknowledgements

We would like to thank Paul Rowley for provision of venoms and Camille Abada and Iara Cardoso for venepuncture. The authors acknowledge the use of the Biomedical Services Unit provided by Liverpool Shared Research Facilities, Faculty of Health and Life Sciences, University of Liverpool. We would also like to thank AstraZeneca Compound Management team in Alderley Park (United Kingdom) for the logistics involving the DMPK studies and Dr. Craig Robertson for assistance with X-ray crystallography analysis. This work was funded by the Wellcome Trust (221712/Z/20/Z to N.R.C. J.K. N.G.B. and P.M.O.). For the purpose of open access, the authors have applied a CC BY public copyright licence to any Author Accepted Manuscript version arising from this submission.

## Conflict of interest

The authors declare no competing interests.

**Correspondence** and requests for material should be addressed to Nicholas R. Casewell and Paul M. O’Neill.

