## Supplementary table 1 for "Discovery and Development of DC-174 as a Novel Oral Snakebite Treatment"

*Supplementary Figures*

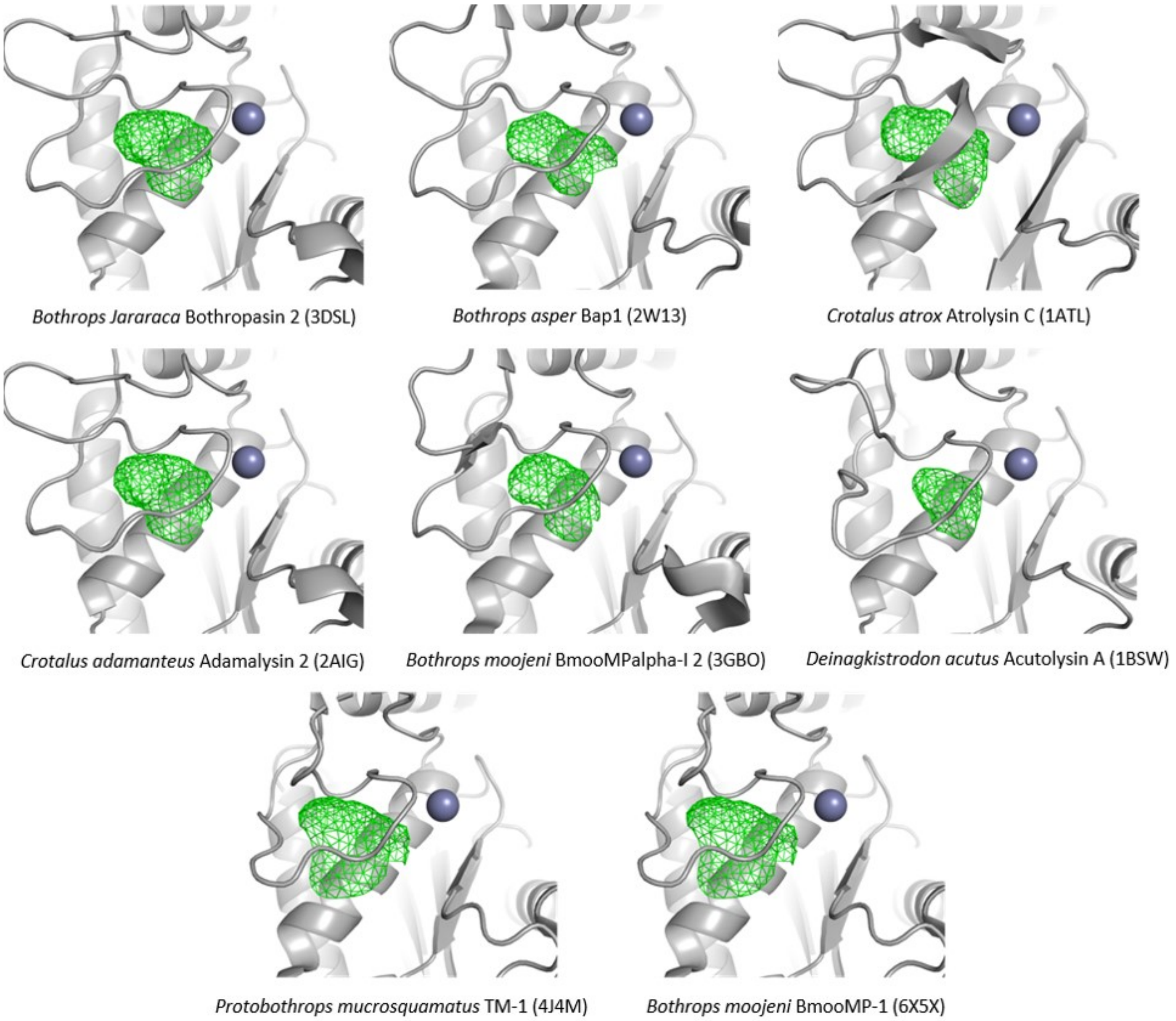


**Supplementary figure 1**. **Structures of SVMPs from different species of snake available in the PDB database.** Accession codes are given in parentheses. Protein secondary structure is represented by grey cartoon, the S1' pocket volume is shown as green wireframe and the catalytic zinc ion is shown as a blue sphere.

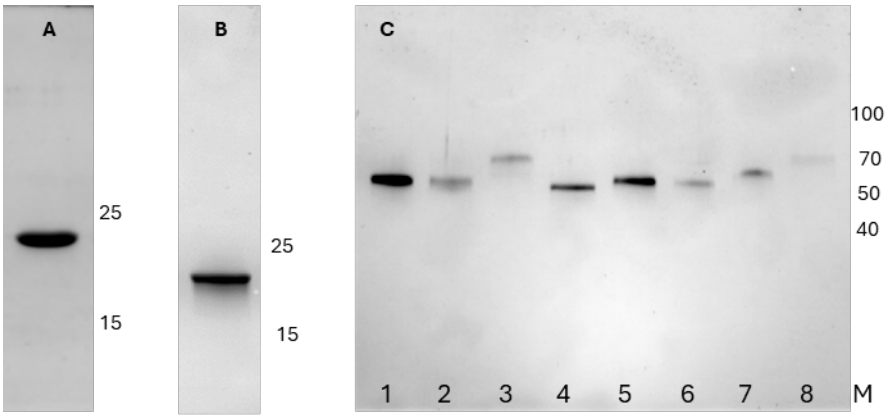


**Supplementary figure 2. SDS-PAGE analysis of the purified *E. romani* SVMPs.** Gel **A**: 22 kDa P-I SVMP. The gel used was 15% acrylamide, made in-house. Gel **B**: 21 kDa P-II SVMP. The gel used was 4-20% acrylamide (BioRad). Gel **C**: P-III SVMPs. The gel used was BioRad 'Any kD'. Lane 1-8, the eight main forms of 52-68 kDa P-III SVMPs individually isolated. When run under non-reducing conditions, the P-III SVMPs in lanes 4 and 5 were shown to be P-IIIc (dimer) forms, the rest were P-IIIa (monomers). All eight were pooled to provide a P-III SVMP sample for use in this study. All gels were stained with Coomassie Blue R250 and key molecular weight markers (Thermo Page Ruler) are indicated in kDa on the right.

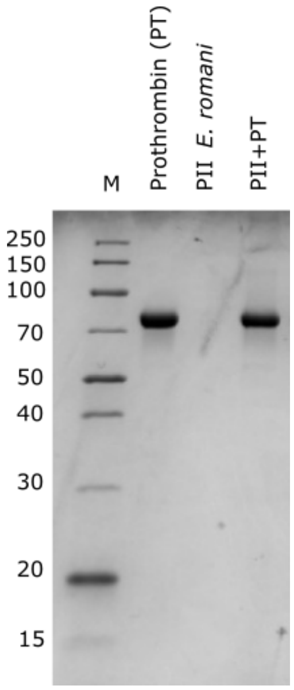


**Supplementary figure 3. The P-II SVMP from *E. romani* venom does not degrade prothrombin.** Prothrombin degradation gel in the presence of P-II SVMP isolated from *E. romani*. Prothrombin (2 µg), SVMP II (200 ng) or prothrombin pre-incubated with 200 ng SVMP II for 1h at 37 ºC (lane 3) were run on an SDS-PAGE gel and stained with Coomassie Brilliant Blue. Protein molecular weights of the marker (M) in kDa are shown on the left.

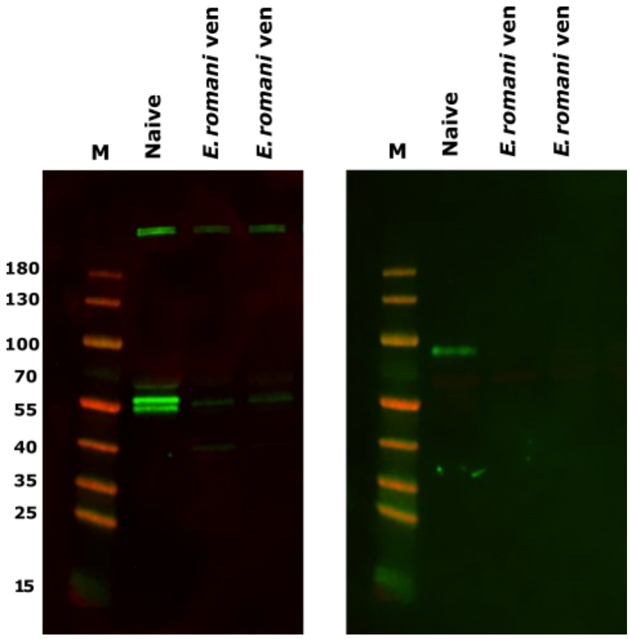


**Supplementary figure 4.** ***E. romani*** **venom depletes fibrinogen and prothrombin** ***in vivo*****.** Anti-fibrinogen (left) and anti-prothrombin (right) western blots of plasma from mice injected intraperitoneally with 90 µg of *E. romani* (n=2) versus naïve controls. Whereas the three chains of murine fibrinogen (alpha, beta and gamma, ~55-65 kDa) are present for the naïve unenvenomed control, a single faint beta chain band remains following the administration of *E. romani* venom (left). Complete degradation of murine prothrombin is seen in envenomed mice (right). M – prestained molecular weight marker.

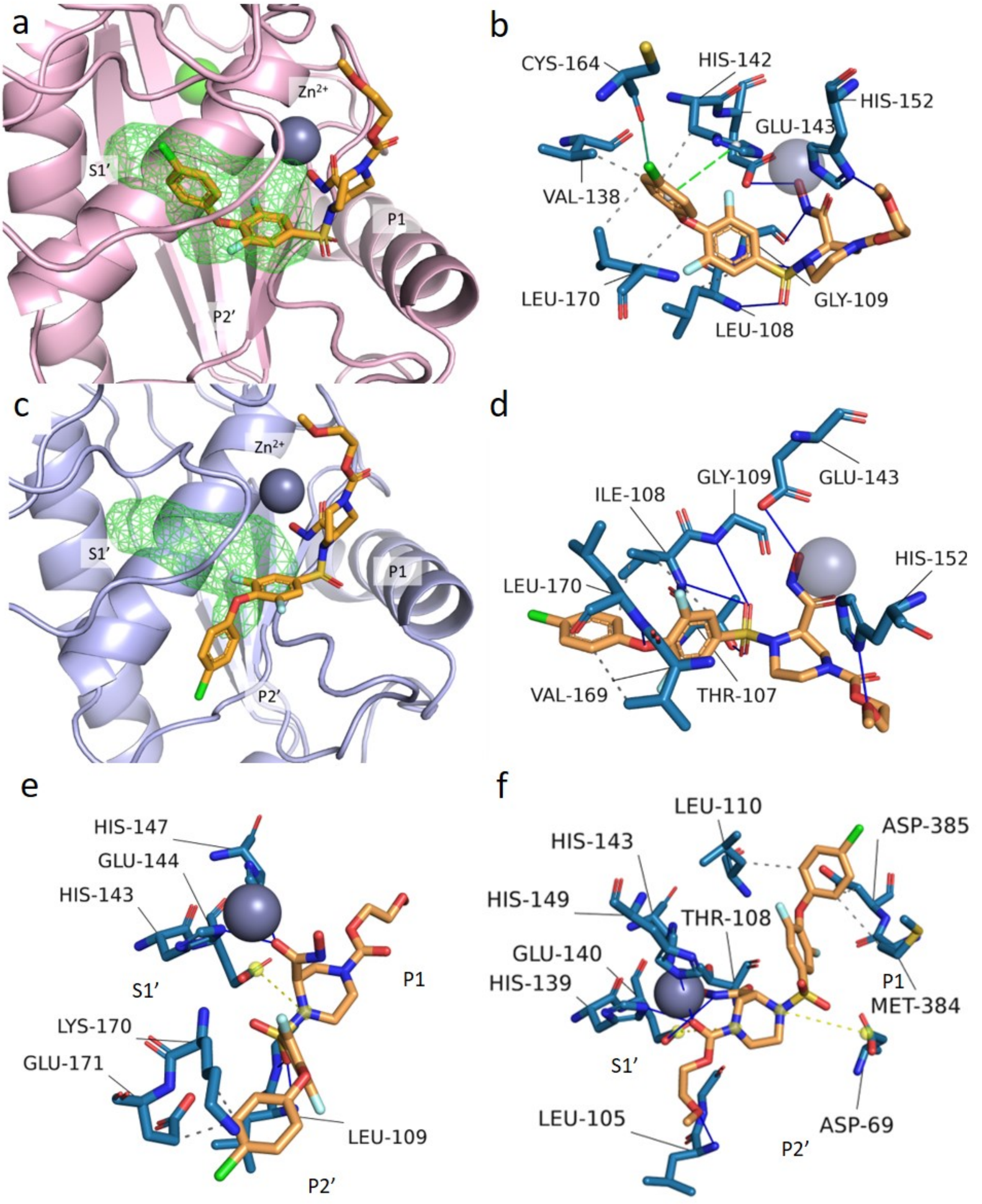


**Supplementary figure 5. Molecular modelling predicted binding poses and interactions between SVMPs and XL-784**. Docked poses of XL-784 (orange sticks) in atrolysin C SVMP (pink cartoons, PDB: 1ATL) (a, b) and BAP-1 SVMP (light blue cartoons, PDB: 2W13) (c, d), ecarin (PDB: 9CLP) (e) and RVV-X (PDB: 2E3X) (f). a, c) Structure of the docked inhibitors in the SVMP binding site, the S1' pocket is highlighted in green wireframe. $Zn^{2+}$ and $Ca^{2+}$ ions are shown in deep blue and green, respectively. b, d, e, f) Analysis of non-covalent interactions formed by inhibitors in the docked structures. Binding residues are shown in blue, hydrophobic interactions are shown as grey dashed lines, salt bridges are shown in yellow dashed lines, π-stacking interaction are shown as green dashed lines, halogen bonds are shown as green solid lines, and hydrogen bonding interactions are shown as blue solid lines.

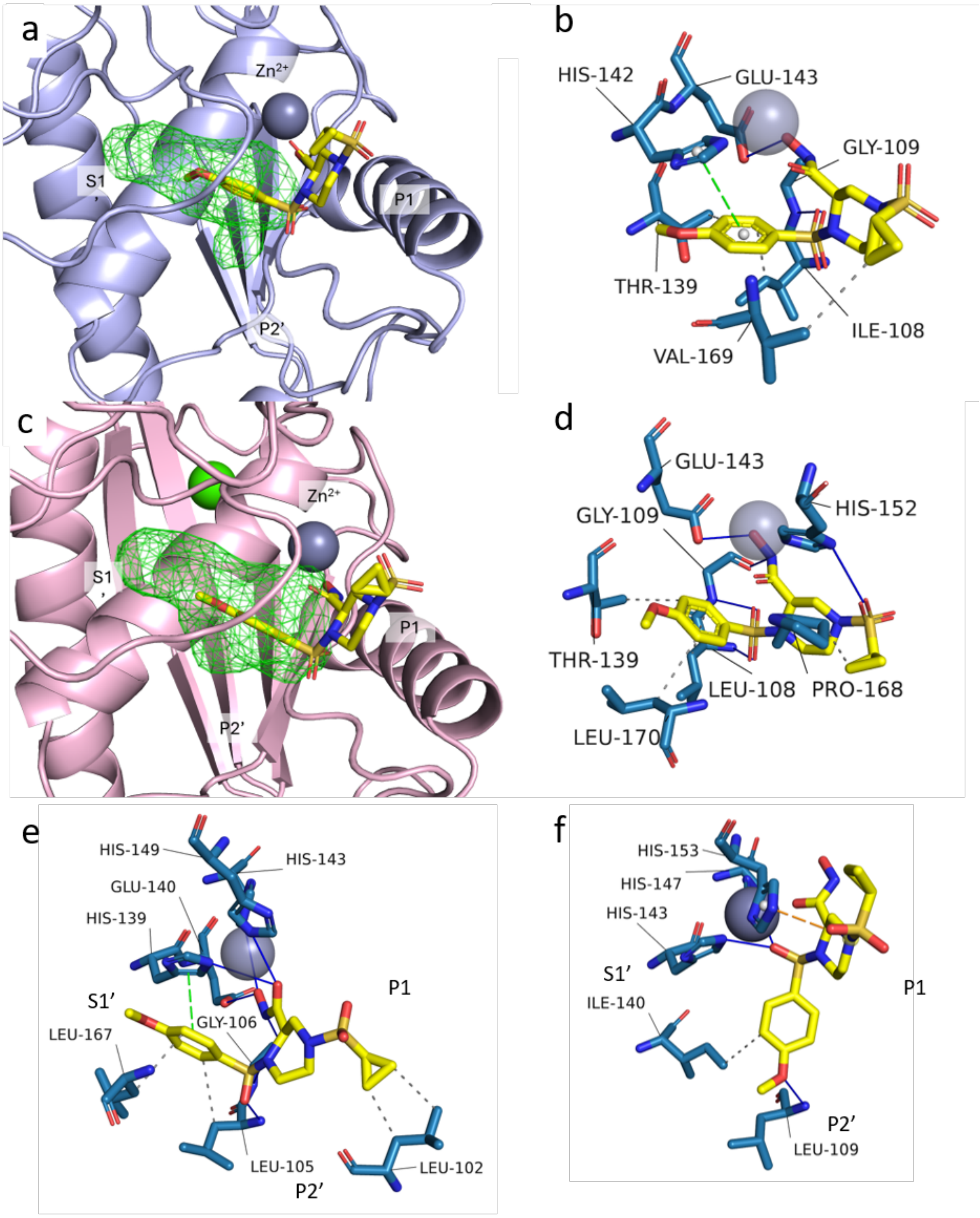


**Supplementary figure 6. Molecular modelling predicted binding poses and interactions between SVMPs and DC-174.** Docked poses of DC-174 (yellow sticks) in BAP-1 SVMP (light blue cartoons, PDB: 2W13) and atrolysin C SVMP (pink cartoons, PDB: 1ATL). a, c) Structure of the docked inhibitors in the SVMP binding site, the S1' pocket is highlighted in green wireframe. $Zn^{2+}$ and $Ca^{2+}$ ions are shown in deep blue and green, respectively. b, d) Analysis of non-covalent interactions formed by inhibitors in the docked structures. Binding residues are shown in blue, hydrophobic interactions are shown as grey dashed lines, π-stacking interaction are shown as green dashed lines, and hydrogen bonding interactions are shown as blue solid lines.

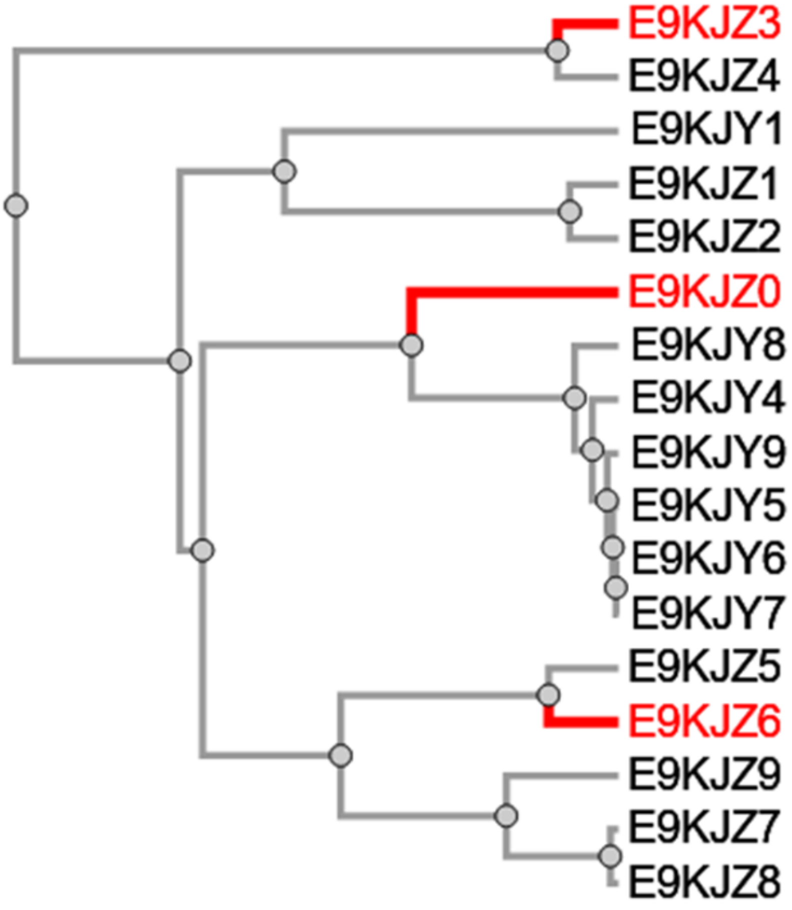


**Supplementary figure 7**. **Phylogenetic tree of P-III sequences denoted by their UniProt identifier.** Identifiers highlighted in red denote the P-III sequences used to build homology models for this work. Primary sequence data for *E. romani* snake venom metalloproteinases (SVMPs) were obtained from UniProt (EoMP06, sequence ID Q6X1T6), comprising two P-I, seven P-II, and 17 P-III SVMP sequences; one representative sequence each from the P-I and P-II subclasses was selected based on sequence coverage, while three P-III sequences were chosen using a phylogenetic analysis conducted with the EMBL-EBI phylogenetics tool in Clustal Omega[5]. Homology models were generated using the SWISS-MODEL web API, model templates were automatically assigned based on sequence similarity[6]. We selected the highest scoring model with the catalytic zinc ion present. Model quality metrics, including Global Model Quality Estimation (GMQE), QMEANDisCo global score, and template coverage, were evaluated, and all selected models passed reliability criteria detailed in Supplementary table S2, which also summarizes UniProt identifiers and corresponding template structures. All models produced had a QMEANDisCo score of > 0.6 indicating reasonable model quality[7].

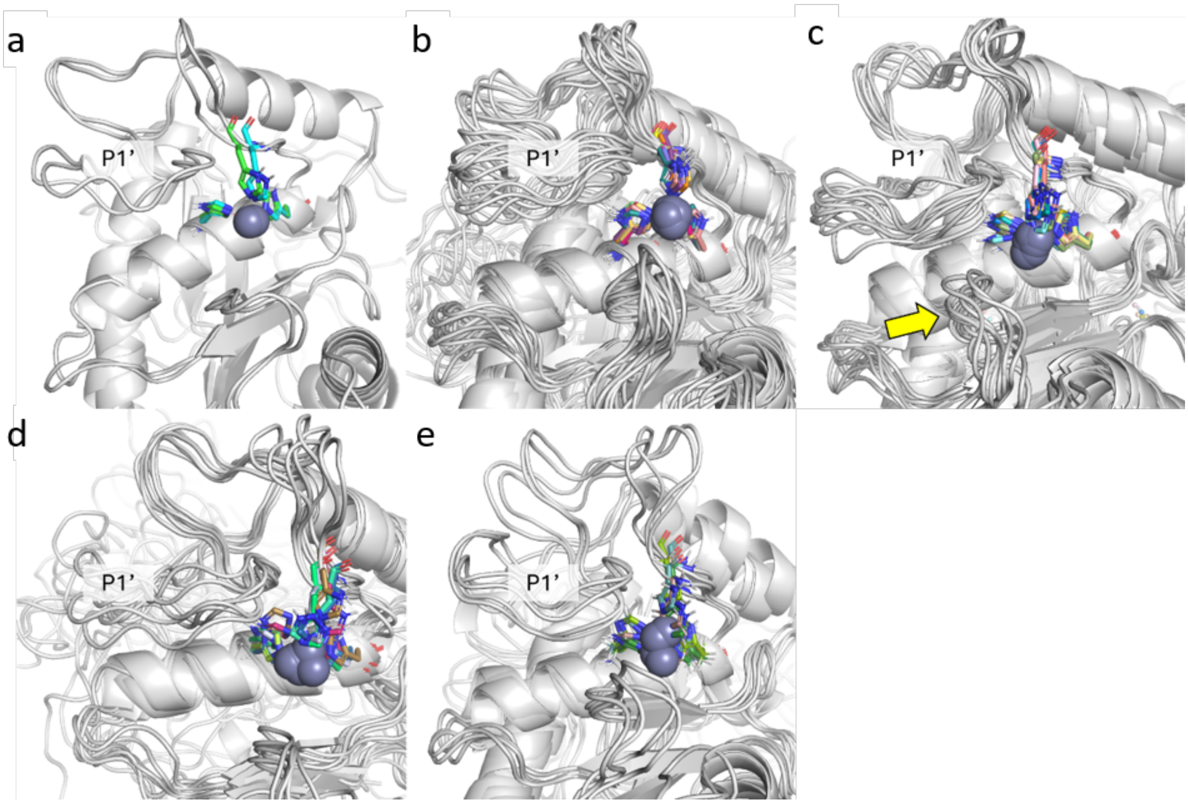


**Supplementary figure 8**. **Conformational ensembles extracted from MD simulations of *E. romani* SVMP subtype homology models**; a) P-I, b) P-II, c) P-III_1, d) P-III_2, e) P-III_3. The P1' pocket is annotated in each panel. In panel c, an adjacent loop which was observed to occlude the P1' pocket is highlighted by the yellow arrow. Ensemble docking was carried out using the CCDC GOLD Python API[8]. Docking was carried out as described as in the main text with modifications to automate setting of harmonic distance constraints using the Gold Python API. The top ten binding poses as determined by ChemPLP scoring function were visually analysed for presence of the ideal binding mode. All scripts used in this work and Pymol sessions of docked structures are available at https://github.com/cmwoodley/SVMP_ensemble_docking.

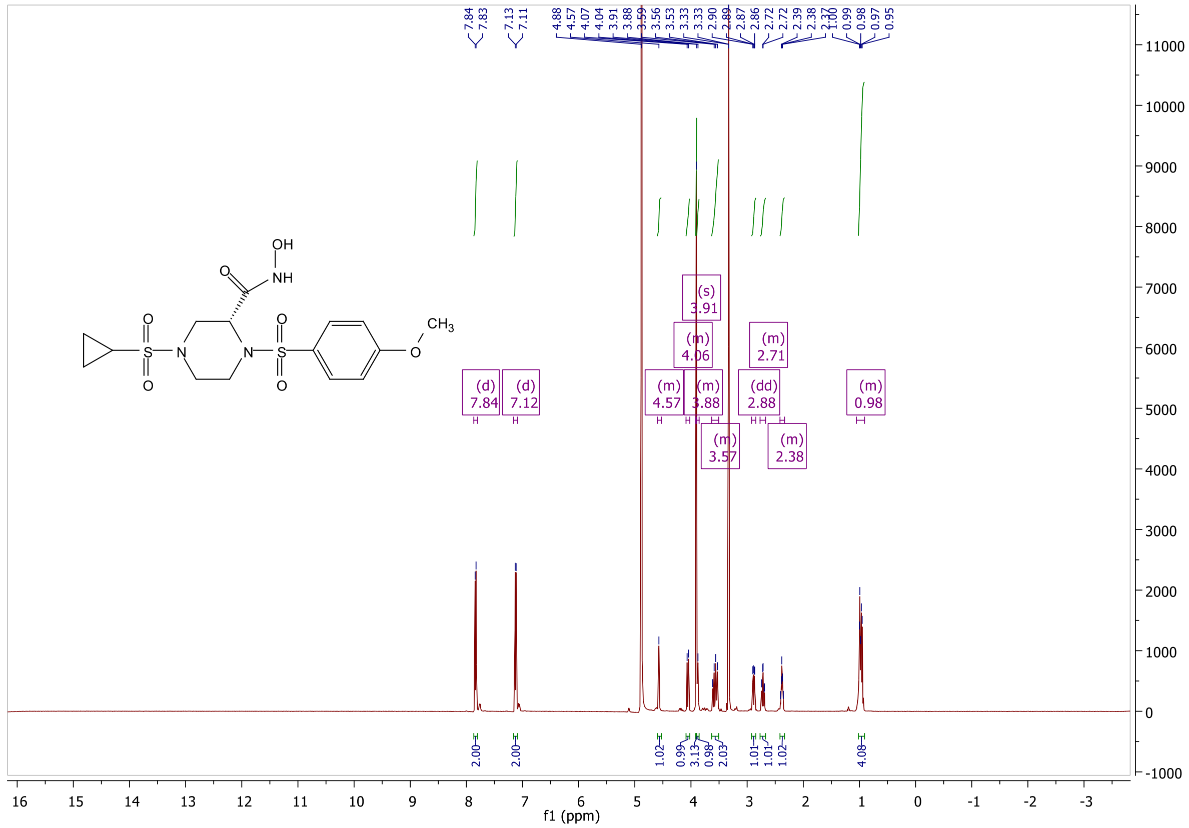

**Supplementary figure 9**. Proton Nuclear Magnetic Resonance spectrum of compound **23** (DC-174).

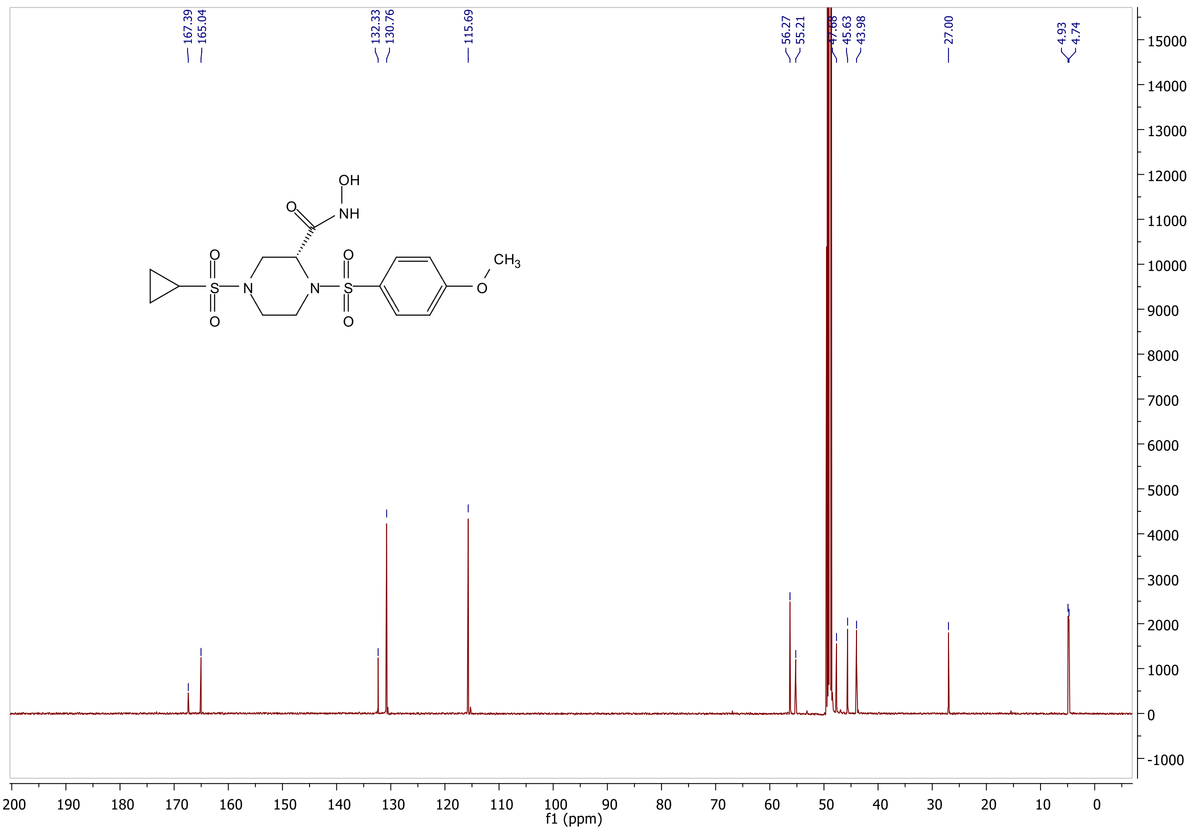


**Supplementary figure 10**. Carbon Nuclear Magnetic Resonance spectrum of compound **23** (DC-174).

### *Supplementary Tables*

**Supplementary table 1**. *In vitro* biological evaluation of compounds described in Figure 2.

| No. | Compound | Inhibition of SVMP enzymatic activity $EC_{50}$ (nM) | | | | | DMPK | | | | |
|---|---|---|---|---|---|---|---|---|---|---|---|
| | | ERO | BJA | BAR | CAT | CRH | $LogD_{7.4}$ | Solubility pH 7.4 (μM) | Human Protein binding (%free) | Rat Heps Met Clint (μL/min/$10^6$ cells) | Hu Mics Metab Clint (μL/min/mg) |
| XL-784 | 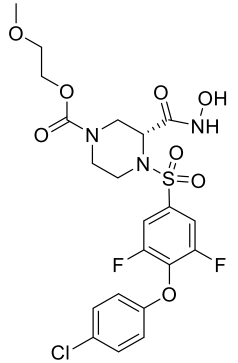 | 9.7 (9.3 - 10.1) | 64.4 (62.1 - 66.7) | 12.9 (12.2 - 13.6) | 50.6 (47.7 - 53.6) | 93.4 (89.7 - 97.1) | 3.2 | 316 | 0.3 | 67.3 | 100.0 |
| **15** | 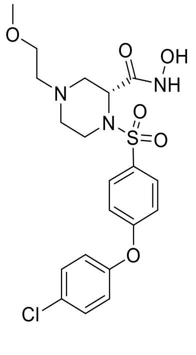 | 7.9 (7.0 - 8.8) | 232.0 (205.7 - 258.3) | 19.9 (18.4 - 21.5) | 47.9 (42.1 - 53.8) | 22.5 (18.9 - 26.2) | 2.4 | 431 | 2.3 | >300.0 | <3.0 |

| | | | | | | | | | | | |
|---|---|---|---|---|---|---|---|---|---|---|---|
| **16** | 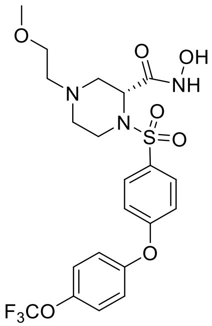 | 15.9 (14.0 - 17.8) | 856.1 (790.4 - 921.7) | 27.8 (22.1 - 33.6) | 76.9 (60.6 - 93.2) | 79.3 (73.3 - 85.3) | 2.9 | 437 | 3.0 | 65.6 | 8.0 |
| **17** | 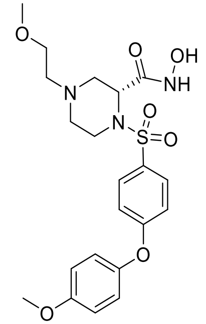 | 1.1 (0.7 - 1.5) | 209.5 (146.0 - 273.0) | 10.4 (4.1 - 16.7) | 31.2 (14.9 - 47.4) | 11.9 (7.7 - 16.1) | 1.9 | 236 | 6.4 | 209.0 | >300 |
| **28** | 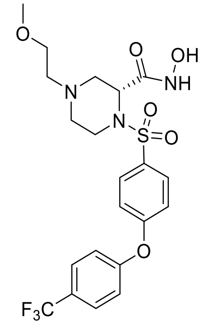 | 7.6 (6.9 - 8.2) | 1512.8 (1435. 2 - 1590.5 ) | 39.5 (36.2 - 42.8) | 68.8 (62.5 - 75.1) | 95.9 (87.5 - 104.3) | 2.7 | 320 | 3.4 | 22.8 | 8.6 |

| | | | | | | | | | | | |
|---|---|---|---|---|---|---|---|---|---|---|---|
| **29** | 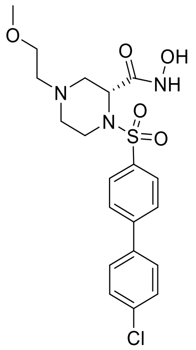 | 16.1 (14.6 - 17.5) | 38.9 (36.2 - 41.7) | 3.5 (3.2 - 3.9) | 52.7 (48.4 - 57.1) | 220.2 (206.8 - 233.7) | 2.5 | 310 | 5.2 | 21.1 | 5.1 |
| **30** | 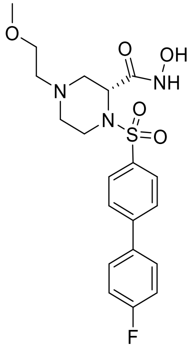 | 18.5 (17.5 - 19.4) | 27.1 (25.6 - 28.6) | 1.8 (1.5 - 2.1) | 18.6 (17.3 - 19.8) | 124.8 (114.8 - 134.8) | 1.9 | 468 | 14.0 | >300 | 7.1 |
| **18** | 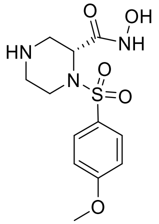 | 47.6 (40.7 - 54.5) | 56.1 (50.9 - 61.2) | 24.7 (21.7 - 27.8) | 69.3 (64.8 - 73.8) | 83.3 (77.2 - 89.4) | -0.2 | 231 | 91 | 6.3 | <3.0 |
| **19** | 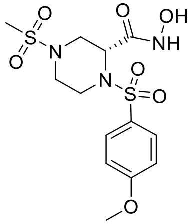 | 10.9 (10.1 - 11.7) | 20.0 (18.6 - 21.5) | 14.3 (13.5 - 15.1) | 45.3 (43.4 - 47.1) | 62.9 (60.1 - 65.6) | -0.3 | <1000 | 88 | 8.9 | <3.0 |

| | | | | | | | | | | | |
|---|---|---|---|---|---|---|---|---|---|---|---|
| **20** | 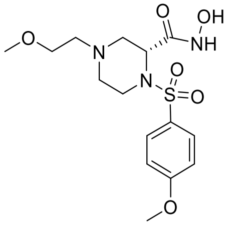 | 61.9 (55.7 - 68.1) | 86.2 (76.4 - 96.1) | 11.1 (9.2 - 13.0) | 68.9 (63.0 - 74.7) | 137.4 (129.3 - 145.4) | 0.1 | >1000 | 82 | 57.1 | <3.0 |
| **21** | 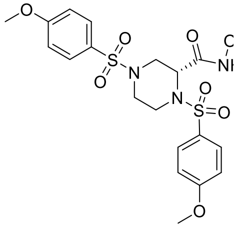 | 15.0 (14.6 - 15.5) | 18.2 (17.4 - 19.0) | 10.6 (10.1 - 11.2) | 32.5 (31.6 - 33.4) | 58.8 (57.0 - 60.6) | 1.0 | 25 | 30 | 199.0 | 10.2 |
| **22** | 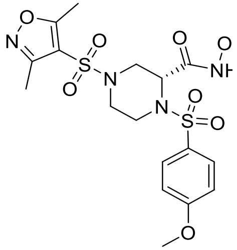 | 10.7 (10.3 - 11.1) | 17.8 (17.1 - 18.6) | 9.8 (9.2 - 10.3) | 35.0 (34.2 - 35.8) | 53.3 (51.8 - 54.8) | 0.7 | >1000 | 51 | 44.0 | 17.5 |
| **23** (DC-174) | 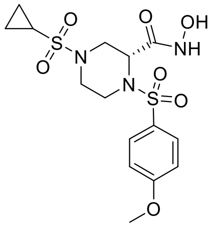 | 4.7 (4.2 - 5.1) | 10.9 (10.2 - 11.7) | 3.9 (3.5 - 4.3) | 22.9 (21.6 - 24.2) | 38.6 (38.3 - 38.8) | 0.2 | >1000 | 66 | 13.3 | 8.3 |

ERO, *E. romani*; BJA, *B. jararaca*; BAR, *B. arietans*; CAT, *C. atrox*; CRH, *C. rhodostoma*.
$LogD_{7.4}$ – Refers to the distribution coefficient between organic and aqueous solvent at physiological pH of 7.4.
Rat Heps Met Clint – Refers to the rate of *in vitro* hepatic intrinsic clearance in rat hepatocytes.
Hu Mics Metab Clint – Refers to the rate of *in vitro* hepatic intrinsic clearance in human microsomes.
$EC_{50}$ data represents means of n= ≥3 repeats, bracketed values represent the range across repeats.

**Supplementary table 2**. The determination of absolute configuration of DC-174 using X-ray crystallography analysis (CCDC deposition number: 2451930).

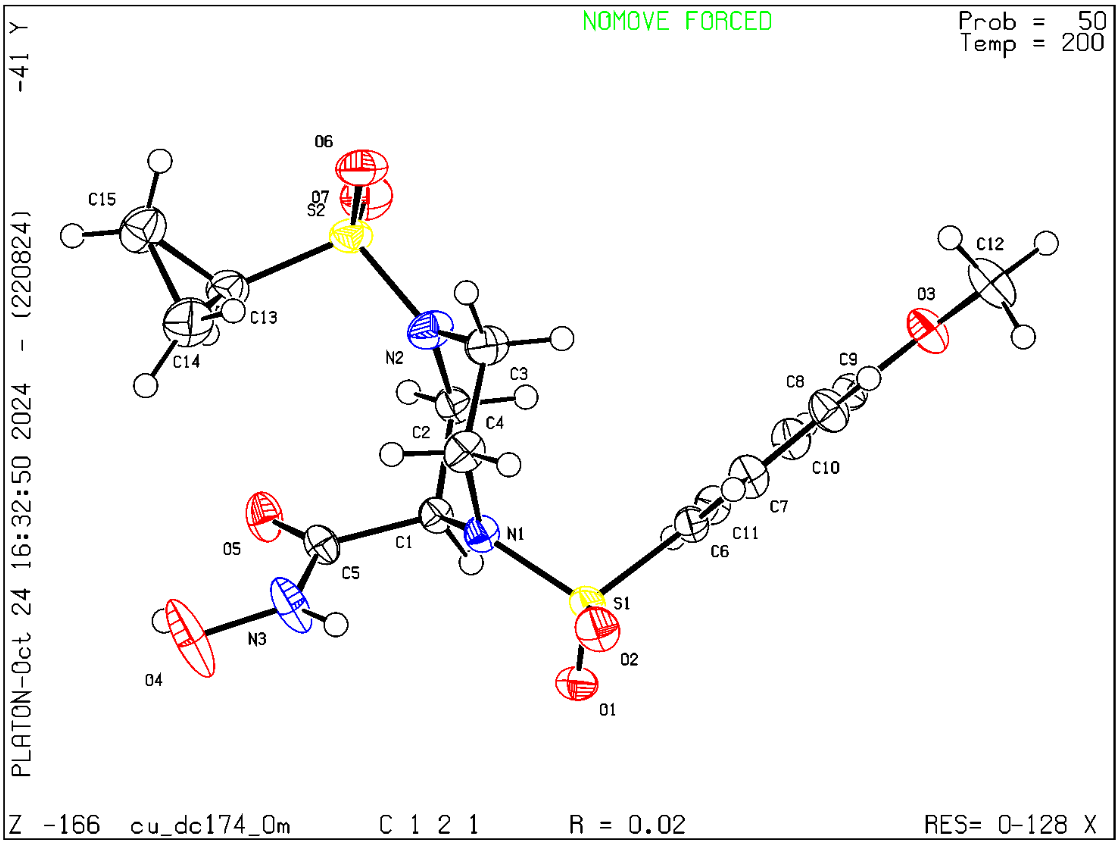


Crystal data and structure refinement for DC-174.

| | |
|---|---|
| Identification code | cu_DC174_0m |
| Empirical formula | $C_{15}H_{21}N_3O_7S_2$ |
| Formula weight | 419.47 |
| Temperature/K | 200.0 |
| Crystal system | monoclinic |
| Space group | C2 |
| a/Å | 19.7656(5) |
| b/Å | 7.4345(2) |
| c/Å | 14.2819(3) |
| α/° | 90 |
| β/° | 118.6210(10) |
| γ/° | 90 |
| Volume/Å$^3$ | 1842.24(8) |
| Z | 4 |
| $\rho_{calc}$g/cm$^3$ | 1.512 |
| μ/mm$^{-1}$ | 3.029 |
| F(000) | 880.0 |

| | |
|---|---|
| Crystal size/mm$^3$ | 0.45 × 0.225 × 0.07 |
| Radiation | CuKα (λ = 1.54178) |
| 2Θ range for data collection/° | 7.05 to 149.34 |
| Index ranges | $-24 \le h \le 24$, $-8 \le k \le 9$, $-17 \le l \le 17$ |
| Reflections collected | 20717 |
| Independent reflections | 3595 [$R_{int}$ = 0.0284, $R_{sigma}$ = 0.0272] |
| Data/restraints/parameters | 3595/1/249 |
| Goodness-of-fit on $F^2$ | 1.092 |
| Final R indexes [I>=2σ (I)] | $R_1$ = 0.0244, $wR_2$ = 0.0637 |
| Final R indexes [all data] | $R_1$ = 0.0245, $wR_2$ = 0.0638 |
| Largest diff. peak/hole / e Å$^{-3}$ | 0.17/-0.31 |
| Flack parameter | 0.057(4) |

**Supplementary table 3**. *In vitro* coagulation assay of selected compounds.

| Selected Inhibitors | Coagulation assay $EC_{50}$ values (nM) | |
|---|---|---|
| | *E. romani* | *B. jararaca* |
| Prinomastat | 557.9 (490.7 - 625.1) | 971.9 (892.2 - 1051.6) |
| XL-784 | >10000 | >10000 |
| **29** | >10000 | >10000 |
| **30** | >10000 | >10000 |
| **19** | 495.6 (447.4 - 543.8) | 509.3 (421.7 - 596.8) |
| **20** | >10000 | >10000 |
| **21** | 4586.6 (3786.6 - 5386.5) | 3919.4 (3324.1 - 4514.6) |
| **22** | 4852.0 (4220.3 - 5483.7) | 1094.9 (926.0 - 1263.8) |
| **23** (DC-174) | 135.0 (114.4 - 155.7) | 13.4 (10.2 - 16.6) |

$EC_{50}$ data represents means of n= ≥3 repeats with range represented in brackets.

**Supplementary table 4.** Summary of protein sequences used for homology model building

| **Model** | **Code** | **Uniprot ID** | **Sequence Description** [a] | **Template** [b] | **Template Desription** | **GMQE** [c] | **QMEANDisCo** [d] |
|---|---|---|---|---|---|---|---|
| 1 | PI | E9KJX3 | Group I snake venom metalloproteinase | 2DW2 | VAP2 from *Crotalus atrox* venom | 0.43 | 0.68 |
| 2 | PII | E9KJX0 | Group II snake venom metalloproteinase | 2DW2 | VAP2 from *Crotalus atrox* venom | 0.45 | 0.68 |
| 3 | PIII_1 | E9KJZ0 | Group III snake venom metalloproteinase | 2DW2 | VAP2 from *Crotalus atrox* venom | 0.62 | 0.74 |
| 4 | PIII_2 | E9KJZ3 | Group III snake venom metalloproteinase | 2DW0 | VAP2 from *Crotalus atrox* venom | 0.79 | 0.72 |
| 5 | PIII_3 | E9KJZ6 | Group III snake venom metalloproteinase | 2DW0 | VAP2 from *Crotalus atrox* venom | 0.62 | 0.74 |

[a] All sequences are from *E. romani* [b] PDB accession codes of templates used to build the homology model [c] GMQE – Global Model Quality stimation [d] QMEANDisCo – Quantitative Model Energy Analysis with Distance Contstraints

**Supplementary table 5**. Summary of ensemble docking into *E. romani* SVMP subtypes

| | | **23** | | | | Prinomastat | | | | XL-784 | | | |
|---|---|---|---|---|---|---|---|---|---|---|---|---|---|
| Code | Interaction | Top 1[a] | Top 3[b] | Top 10[c] | Ideal[d] | Top 1[a] | Top 3[b] | Top 10[c] | Ideal[d] | Top 1[a] | Top 3[b] | Top 10[c] | Ideal[d] |
| P-I | Binds P1’ | Y | 3/3 | 10/10 | 5/10 | Y | 3/3 | 10/10 | 7/10 | Y | 3/3 | 9/10 | 4/10 |
| | Sulfone HB | N | 1/3 | 5/10 | | Y | 3/3 | 7/10 | | N | 1/3 | 5/10 | |
| P-II | Binds P1’ | N | 1/3 | 4/10 | 3/10 | Y | 1/3 | 5/10 | 2/10 | N | 0/3 | 0/10 | 0/10 |
| | Sulfone HB | Y | 2/3 | 5/10 | | N | 0/3 | 2/10 | | N | 1/3 | 1/10 | |
| P-III_1 | Binds P1’ | N | 0/3 | 0/10 | 0/10 | Y | 0/3 | 0/10 | 0/10 | N | 0/3 | 0/10 | 0/10 |
| | Sulfone HB | N | 0/3 | 0/10 | | N | 0/3 | 0/10 | | N | 0/3 | 0/10 | |
| P-III_2 | Binds P1’ | N | 0/3 | 2/10 | 1/10 | N | 0/3 | 2/10 | 1/10 | N | 0/3 | 0/10 | 0/10 |
| | Sulfone HB | N | 0/3 | 2/10 | | N | 0/3 | 1/10 | | N | 0/3 | 1/10 | |
| P-III_3 | Binds P1’ | Y | 3/3 | 9/10 | 5/10 | Y | 3/3 | 10/10 | 1/10 | Y | 1/3 | 5/10 | 4/10 |
| | Sulfone HB | Y | 2/3 | 5/10 | | N | 1/3 | 1/10 | | Y | 1/3 | 4/10 | |
| | **Total P1’:** | 2 | 7 | 25 | | 4 | 7 | 27 | | 2 | 4 | 14 | |
| | **Total Sulfone:** | 2 | 5 | 17 | | 1 | 4 | 11 | | 1 | 3 | 11 | |
| | **Totals:** | 4 | 12 | 42 | 14 | 5 | 11 | 38 | 11 | 3 | 7 | 25 | 8 |

[a] Presence of a given interaction in the top scoring pose [b] Presence of a given interaction in the top 3 poses [c] Presence of a given interaction in the top 10 poses [d] Presence of the ideal binding mode, defined by a pose in which the aryl substituent is bound within the P1’ pocket and the sulfonamide forms hydrogen binding interactions with the protein backbone.

***Supplementary information for molecular modelling.***

### Additional discussion for molecular docking studies

Docking studies using molecular dynamics-derived ensembles of *E. romani* SVMP homology models (one P-I, one P-II, and three P-III; see Supplementary figure 7 sequence selections for homology modelling; see Supplementary table 4 for of protein sequences used for homology model building; Supplementary table 5 for summary of ensemble docking) were performed to assess binding of DC-174, prinomastat, and XL-784. Both DC-174, and prinomastat formed binding modes across all SVMP subtypes (see Supplementary figure 8). However, XL-784 failed to bind effectively to P-II SVMPs, likely due to its bulky biaryl side chain not fitting in the P1' pocket. These findings align with fibrinogen cleavage data (Figure 3a) and are further detailed in the Supporting Information.

### Molecular modelling of *E. romani* SVMP isoforms

A limitation of docking into available X-ray structures of SVMPs is that it does not account for the diversity of SVMP isoforms within a species. Our gel electrophoresis experiments demonstrated that different molecules exhibited distinct inhibition profiles against various SVMP subtypes from *E. romani*. To address this, we docked DC-174, prinomastat, and XL-784 into homology models constructed from the primary sequences of *E. romani* P-I, P-II, and P-III SVMP subtypes (see Supplementary table 4 for protein sequences used for homology model building). The P1' pocket is known to be flexible[9,10], so to sample conformational space, we performed multiple short unbiased molecular dynamics (MD) simulations using the homology models as starting coordinates. Representative structures were then extracted through clustering of the resulting trajectories[11]; these ensembles are shown in Supplementary figure 8. In the first P-III isoform (Supplementary figure 8c) an adjacent disorganised loop occluded the P1' pocket in each representative structure; in the docking studies this prevented formation of interactions with the P1' pocket and backbone hydrogen bonds with the sulfonamide.

We used the observed binding modes to qualitatively compare ligands based on the number of poses in an ideal binding mode - defined as the aryl group occupying the P1' pocket and the sulfonamide forming hydrogen bonds with the protein backbone. The main limitation of this workflow is that we do not consider the free energy landscape of P1' pocket opening. This limits the inference of this method to assessing whether or not a ligand can form an ideal binding mode, rather than assessing how likely a ligand is to bind. However, due to the biological complexity of SVMPs within a given snake species venom, we believe this is a reasonable compromise considering conformational flexibility for multiple isoforms while balancing computational expense. A summary of these qualitative results of this ensemble docking is provided in Supplementary table 5.

Consistent with docking results using X-ray structures, DC-174 formed ideal binding modes across all *E. romani* SVMP isoforms, with the largest number of ideal binding poses compared to Prinomastat and XL-784. Prinomastat also formed ideal binding modes with all isoforms, while XL-784 failed to produce any ideal binding poses with P-II SVMPs. This aligns with the poor inhibition of fibrinogen cleavage by XL-784 observed in gel electrophoresis assays (Figure 3a), where both DC-174 and Prinomastat showed significant inhibition. In our homology models, we observed the presence of an arginine residue in the P1' pocket of P-II SVMP, which may act as a gatekeeper, similar to the arginine in MMP1 that plays a crucial role in the selective inhibition of MMP13[9,12]. This arginine residue likely prevents the bulky side chain of XL-784 from effectively binding within the P1' pocket of P-II SVMP.

Further analysis of the number of poses with the aryl group bound in the P1' pocket revealed comparable results for DC-174 and Prinomastat, whereas XL-784 showed fewer poses. This suggests that the bulkiness of the biaryl side chain in XL-784 hinders its ability to bind effectively in the P1' pocket, consistent with secondary assay observations. These studies highlight that truncating the aryl substituent to a single ring may be a viable strategy to achieve broad-spectrum activity both within and across species.

**Molecular dynamics**

To sample the flexibility of the P1 pocket, we used each homology model as starting coordinates for molecular dynamics simulations. For each SVMP model we carried out five replica 5 ns simulations, generating new velocities for each replicate run. The molecular dynamics (MD) simulations were performed using Gromacs 2022.0 and the Amber FF99-SB ILDN force field[13]. Following the preparation of the system, steepest descent energy minimization was conducted for 50,000 steps. The system was subsequently equilibrated for 100 ns in the NVT followed by 100 ns simulation in the NPT ensemble with positional restraints on protein heavy atoms. The production MD was performed with a leap-frog integrator using a time step of 2 fs for a total simulation duration of 5 ns. Long-range electrostatics were computed using the Particle Mesh Ewald (PME) method, with a real-space cut off of 1.2 nm, and van der Waals interactions were treated with a force-switch modifier within a 1.0–1.2 nm cut off. LINCS constraints were applied to all bonds involving hydrogen. Temperature was controlled using a modified Berendsen thermostat (V-rescale) at 300 K[14], with separate coupling groups for the protein and solvent, and pressure was maintained at 1 bar using the Parrinello-Rahman barostat[15]. Output coordinates, energies, and logs were saved every 10 ps.

Mdtraj was used to process trajectories[16]. Trajectories were pre-processed by discarding the first nanosecond of each for equilibration – totalling 20 ns simulation time for each subtype. Trajectories were concatenated and clustered into 12 clusters each using agglomerative clustering as implemented in scikit-learn[17] based on the root-mean-square deviation of protein heavy atoms within 12.0 Å of the catalytic zinc ion. Cluster centroids were extracted for use in ensemble docking. In simulations involving P-I, P-III_1, P-III_2 and P-III_3 the catalytic zinc was observed to dissociate – cluster centroids where the zinc had dissociated were not included for ensemble docking. Structures of cluster centroids were aligned to each other based on protein heavy atoms within 12.0 Å of the catalytic zinc using the Pymol python API[18].

***Supplementary information for chemical synthesis procedures.***

**General Procedure for the synthesis of sulfonamides 2-5, 10, 12-14 and 24.**

1-(*tert*-Butyl) 3-methyl (*R*)-piperazine-1,3-dicarboxylate (SM) and 4-dimethylaminopyridine (DMAP) was charged into a 100 mL round bottom flask, sealed with a septum, evacuated and back-filled with nitrogen (2 times). 1,4-Dioxane and triethylamine is added and the reaction mixture was stirred. Appropriate sulfonyl chloride (dissolved in 1,4-dioxane) was added drop-wise and the reaction mixture was stirred for 16 h. Water was added and the aqueous solution was extracted with ethyl acetate (2 x 50 mL). The pooled organic solution was dried with $MgSO_4$, filtered and concentrated *in-vacuo*. The crude product was purified by column chromatography.

*1-(tert-Butyl) 3-methyl (R)-4-((4-(4-chlorophenoxy)phenyl)sulfonyl)piperazine-1,3-dicarboxylate* (**2**). SM (1.00 g, 4.09 mmol), DMAP (50 mg, 0.41 mmol), 1,4-dioxane (30 mL), triethylamine (1.71 mL, 12.28 mmol), 4-(4-chlorophenoxy)benzenesulfonyl chloride (1.24 g, 4.09 mmol, in 5 mL of 1,4-dioxane) following column elution from 0% to 20% ethyl acetate in hexane to give **2** (1.73 g, 3.38 mmol, 83%) as colourless liquid. $^1$H NMR (400 MHz, $CDCl_3$) δ $^1$H NMR (400 MHz, $CDCl_3$) δ 7.74 - 7.69 (d, *J* = 8.8 Hz, 2H), 7.39 – 7.33 (d, *J* = 8.8 Hz, 2H), 7.07 – 6.94 (m, 4H), 4.62 - 4.56 (m, 2H), 4.56 – 4.51 (m, 1H), 4.22 – 3.92 (m, 1H), 3.69 – 3.59 (m, 1H), 3.56 (s, 3H), 3.50 – 3.30 (m, 1H), 3.23 – 3.04 (m, 1H), 3.03 – 2.77 (m, 1H), 1.40 (s, 9H).

*1-(tert-Butyl) 3-methyl (R)-4-((4-(4-(trifluoromethoxy)phenoxy)phenyl)sulfonyl)piperazine-1,3-dicarboxylate* (**3**). SM (128 mg, 0.52 mmol), DMAP (7 mg, 0.057 mmol), 1,4-dioxane (5 mL), triethylamine (0.15 mL, 1.57 mmol), 4-(4-(trifluoromethoxy)phenoxy)benzenesulfonyl chloride (185 mg, 0.52 mmol, in 2 mL of 1,4-dioxane) following column elution from 0% to 20% ethyl acetate in hexane to give **3** (220 mg, 0.39 mmol, 75%) as pale-yellow liquid. $^1$H NMR (400 MHz, $CDCl_3$) δ 7.78 – 7.70 (d, *J* = 8.8 Hz, 2H), 7.24 (d, *J* = 9.2 Hz, 2H), 7.07 (d, *J* = 9.2 Hz, 2H), 7.04 (d, *J* = 8.8 Hz, 4H), 4.62 – 4.57 (m, 1H), 4.57 – 4.44 (m, 1H), 4.20 - 3.99 (m, 1H), 3.68 – 3.60 (m, 1H), 3.56 (s, 3H), 3.49 – 3.30 (m, 1H), 3.23 – 3.05 (m, 1H), 3.02 – 2.78 (m, 1H), 1.40 (s, 9H).

*1-(tert-Butyl) 3-methyl (R)-4-((4-(4-methoxyphenoxy)phenyl)sulfonyl)piperazine-1,3-dicarboxylate* (**4**). SM (189 mg, 1.23 mmol), DMAP (14 mg, 0.11 mmol), 1,4-dioxane (10 mL), triethylamine (0.49 mL, 3.69 mmol), 4-(4-methoxyphenoxy)benzenesulfonyl chloride (350 mg, 4.32 mmol, in 5 mL of 1,4-dioxane) following column elution from 0% to 20% ethyl acetate in hexane to give **4** (400 mg, 0.79 mmol, 64%) as pale-yellow liquid. $^1$H NMR (400 MHz, $CDCl_3$) δ 7.68 (d, *J* = 8.8 Hz, 2H), 7.04 – 6.88 (m, 6H), 4.61 – 4.55 (m, 1H), 4.55 - 4.43 (m, 1H), 4.20 - 3.93 (m, 1H), 3.83 (s, 3H), 3.66 – 3.58 (m, 1H), 3.55 (s, 3H), 3.49 – 3.30 (m, 1H), 3.21 – 3.04 (m, 1H), 3.01 – 2.78 (m, 1H), 1.40 (s, 9H).

*1-(tert-Butyl) 3-methyl (R)-4-((4-methoxyphenyl)sulfonyl)piperazine-1,3-dicarboxylate* (**5**). SM (1.00 g, 4.09 mmol), DMAP (50 mg, 0.41 mmol), 1,4-dioxane (30 mL), triethylamine (1.71 mL, 12.28 mmol), 4-methoxybenzenesulfonyl chloride (1.25 g, 4.09 mmol, in 10 mL of 1,4-dioxane) following column elution from 0% to 20% ethyl acetate in hexane to give **5** (1.61 g, 3.88 mmol, 95%) as colourless liquid. $^1$H NMR (400 MHz, $CDCl_3$) δ 7.69 (d, *J* = 9.2 Hz, 2H), 6.95 (d, *J* = 9.2 Hz, 4H), 4.60 – 4.53 (m, 1H), 4.54 – 4.39 (m, 1H), 4.20 – 3.93 (m, 1H), 3.86

(s, 3H), 3.67 – 3.57 (m, 1H), 3.53 (s, 3H), 3.47 – 3.29 (m, 1H), 3.20 – 3.02 (m, 1H), 2.99 – 2.75 (m, 1H), 1.39 (s, 9H).

*Methyl (R)-1-((4-methoxyphenyl)sulfonyl)-4-(methylsulfonyl)piperazine-2-carboxylate* (**10**). Boc-protected **5** (467 mg, 1.12 mmol) was first deprotected with TFA (10 eq.) in DCM (15 mL). The crude reaction mixture was carefully neutralised with saturated sodium bicarbonate solution to give crude methyl *(R)*-1-((4-methoxyphenyl)sulfonyl)piperazine-2-carboxylate. Following that, methyl *(R)*-1-((4-methoxyphenyl)sulfonyl)piperazine-2-carboxylate (356 mg, 1.12 mmol), DMAP (14 mg, 0.11 mmol), 1,4-dioxane (10 mL), triethylamine (0.47 mL, 3.39 mmol), methanesulfonyl chloride (92 µL, 1.19 mmol) following column elution from 0% to 50% ethyl acetate in hexane to give **10** (412 mg, 1.05 mmol, 93%) as colourless liquid. $^1$H NMR (400 MHz, $CDCl_3$) δ 7.73 (d, $J$ = 9.2 Hz, 2H), 6.97 (d, $J$ = 9.2 Hz, 2H), 4.85 – 4.75 (m, 1H), 4.25 – 4.17 (m, 1H), 3.87 (s, 3H), 3.81 - 3.67 (m, 2H), 3.62 (s, 3H), 3.40 (m, 1H), 3.02 (m, 1H), 2.84 (m, 1H), 2.78 (s, 3H).

*Methyl (R)-1,4-bis((4-methoxyphenyl)sulfonyl)piperazine-2-carboxylate* (**12**). Boc-protected **5** (421 mg, 1.02 mmol) was first deprotected with TFA (10 eq.) in DCM (15 mL). The crude reaction mixture was carefully neutralised with saturated sodium bicarbonate solution to give crude methyl *(R)*-1-((4-methoxyphenyl)sulfonyl)piperazine-2-carboxylate. Following that, methyl *(R)*-1-((4-methoxyphenyl)sulfonyl)piperazine-2-carboxylate (318 mg, 1.02 mmol), DMAP (12 mg, 0.098 mmol), 1,4-dioxane (10 mL), triethylamine (0.42 mL, 3.03 mmol), 4-methoxybenzenesulfonyl chloride (310 mg, 1.02 mmol) following column elution from 0% to 50% ethyl acetate in hexane to give **12** (384 mg, 0.79 mmol, 78%) as colourless liquid. $^1$H NMR (400 MHz, $CDCl_3$) δ 7.67 (d, $J$ = 9.2 Hz, 2H), 7.64 (d, $J$ = 9.2 Hz, 2H), 6.99 (d, $J$ = 8.8 Hz, 2H), 6.93 (d, $J$ = 8.8 Hz, 2H), 4.75 – 4.70 (m, 1H), 4.19 – 4.13 (m, 1H), 3.88 (s, 3H), 3.85 (s, 3H), 3.74 – 3.62 (m, 2H), 3.62 (s, 3H), 3.47 – 3.37 (m, 1H), 2.53 (dd, $J$ = 11.7, 4.0 Hz, 1H), 2.32 – 2.30 (m, 1H).

*Methyl (R)-4-((3,5-dimethylisoxazol-4-yl)sulfonyl)-1-((4-methoxyphenyl)sulfonyl)piperazine-2-carboxylate* (**13**). Boc-protected **5** (696 mg, 1.68 mmol) was first deprotected with TFA (10 eq.) in DCM (10 mL). The crude reaction mixture was carefully neutralised with saturated sodium bicarbonate solution to give crude methyl (R)-1-((4-methoxyphenyl)sulfonyl)piperazine-2-carboxylate. Following that, methyl (R)-1-((4-methoxyphenyl)sulfonyl)piperazine-2-carboxylate (528 mg, 2.16 mmol), DMAP (26 mg, 0.21 mmol), 1,4-dioxane (15 mL), triethylamine (0.90 mL, 6.48 mmol), 3,5-dimethylisoxazole-4-sulfonyl chloride (423 mg, 2.16 mmol) following column elution from 0% to 50% ethyl acetate in hexane to give **13** (561 mg, 1.18 mmol, 55%) as yellow liquid. $^1$H NMR (400 MHz, $CDCl_3$) δ 7.70 (d, $J$ = 9.2 Hz, 2H), 6.96 (d, $J$ = 9.2 Hz, 2H), 4.81 – 4.74 (m, 1H), 4.10 – 4.03 (m, 1H), 3.87 (s, 3H), 3.82 – 3.74 (m, 1H), 3.72 – 3.64 (m, 1H), 3.57 (s, 3H), 3.40 (td, $J$ = 12.0, 3.2 Hz, 1H), 2.94-2.85 (m, 1H), 2.72 (td, $J$ = 12.0, 3.2 Hz, 1H), 2.62 (s, 3H), 2.32 (s, 3H).

*Methyl (R)-4-(cyclopropylsulfonyl)-1-((4-methoxyphenyl)sulfonyl)piperazine-2-carboxylate* (**14**). Boc-protected **5** (466 mg, 1.12 mmol) was first deprotected with TFA (10 eq.) in DCM (10 mL). The crude reaction mixture was carefully neutralised with saturated sodium bicarbonate solution to give crude methyl (R)-1-((4-methoxyphenyl)sulfonyl)piperazine-2-carboxylate. Following that, methyl (R)-1-((4-methoxyphenyl)sulfonyl)piperazine-2-carboxylate (354 mg, 1.45 mmol), DMAP (18 mg, 0.15 mmol), 1,4-dioxane (10 mL), triethylamine (0.61 mL, 4.35 mmol), cyclopropanesulfonyl chloride (148 µL, 1.45 mmol) following column elution from 0% to 50% ethyl acetate in hexane to give **14** (365 mg, 0.87 mmol, 60%) as colourless liquid. $^1$H NMR $^1$H NMR (400 MHz, $CDCl_3$) δ 7.73 (d, *J* = 9.2 Hz, 2H), 6.97 (d, *J* = 9.2 Hz, 2H), 4.80 - 4.75 (m, 1H), 4.26 – 4.17 (m, 1H), 3.87 (s, 3H), 3.81 – 3.66 (m, 2H), 3.62 (s, 3H), 3.41 (td, *J* = 12.4, 3.4 Hz, 1H), 3.17 – 3.10 (m, 1H), 2.96 (td, *J* = 12.4, 3.4 Hz, 1H), 2.26 – 2.15 (m, 1H), 1.19 – 1.10 (m, 2H), 1.05 – 0.94 (m, 2H).

*1-(tert-Butyl) 3-methyl (R)-4-((4-bromophenyl)sulfonyl)piperazine-1,3-dicarboxylate* (**24**). SM (1.00 g, 4.09 mmol), DMAP (50 mg, 0.41 mmol), 1,4-dioxane (30 mL), triethylamine (1.71 mL, 12.28 mmol), 4-bromobenzenesulfonyl chloride (1.05 g, 4.09 mmol, in 5 mL of 1,4-dioxane) following column elution from 0% to 20% ethyl acetate in hexane to give **24** (1.34 g, 2.89 mmol, 70%) as pale-yellow liquid. $^1$H NMR (400 MHz, $CDCl_3$) δ 7.67 – 7.58 (m, 4H), 4.62 – 4.57 (m, 2H), 4.57 - 4.45 (m, 1H), 4.24 - 3.93 (m, 1H), 3.70 – 3.59 (m, 1H), 3.54 (s, 3H), 3.47 – 3.28 (m, 1H), 3.21 – 3.04 (m, 1H), 3.02 – 2.75 (m, 1H), 1.40 (s, 9H).

**General Procedure for the alkylation of *N*-Boc piperazine 6-8, 11 and 25.**

In a 50 mL round bottom flask, N-Boc piperazine was treated with trifluoroacetic acid (TFA) in DCM and the reaction mixture was stirred for 16 h. The reaction mixture was concentrated and re-dissolved in DCM. The organic solution was washed carefully with saturated solution of sodium bicarbonate, dried with $MgSO_4$, filtered and concentrated *in-vacuo*. To the resulting crude mass, potassium carbonate is added and the round bottom flask was sealed with a septum. Anhydrous DMF was added and the reaction mixture was stirred for 30 mins. 2-Bromoethyl methyl ether was added drop-wise and the reaction mixture was stirred for 30 mins before being heated to 60°C for 16 h. The reaction mixture was concentrated *in-vacuo* and water was added. The aqueous solution was extracted with ethyl acetate (3 times). The pooled organic solution was dried with $MgSO_4$, filtered and concentrated *in-vacuo*. The crude product was purified by column chromatography.

*Methyl (R)-1-((4-(4-chlorophenoxy)phenyl)sulfonyl)-4-(2-methoxyethyl)piperazine-2-carboxylate* (**6**). **2** (0.79 g, 1.54 mmol), TFA (1.18 mL, 15.40 mmol), DCM (10 mL) to give crude mass of 634 mg. Potassium carbonate (426 mg, 3.08 mmol), DMF (6 mL), 2-bromoethyl methyl ether (290 μL, 3.08 mmol) following column elution from 10% to 60% ethyl acetate in hexane to give **6** (512 mg, 1.09 mmol, 71%) as colourless liquid. $^1$H NMR (400 MHz, $CDCl_3$) δ 7.75 (d, *J* = 8.8 Hz, 2H), 7.35 (d, *J* = 8.8 Hz, 2H), 7.06 – 6.96 (m, 4H), 4.64 – 4.59 (m, 1H), 3.61 (s, 3H), 3.48 – 3.33 (m, 4H), 3.30 (s, 3H), 2.81 – 2.74 (m, 1H), 2.63 – 2.45 (m, 2H), 2.42 – 2.32 (m, 1H), 2.26 (td, *J* = 11.5, 3.6 Hz, 1H).

*Methyl (R)-1-((4-bromophenyl)sulfonyl)-4-(2-methoxyethyl)piperazine-2-carboxylate* (2**5**). **3** (750 mg, 1.85 mmol), TFA (1.42 mL, 18.50 mmol), DCM (10 mL) to give crude mass of 606 mg. Potassium carbonate (461 mg, 3.34 mmol), DMF (6 mL), 2-bromoethyl methyl ether (314 μL, 2.27 mmol) following column elution from 40% to 100% ethyl acetate in hexane to give 2**5** (393 mg, 0.93 mmol, 50%) as colourless liquid. $^1$H NMR (400 MHz, $CDCl_3$) δ 7.68 – 7.60 (m, 4H), 4.64 – 4.60 (m, 1H), 3.65 – 3.59 (m, 1H), 3.58 (s, 3H), 3.46 – 3.31 (m, 4H), 3.29 (s, 3H), 2.83 – 2.74 (m, 1H), 2.63 – 2.46 (m, 2H), 2.37 (dd, *J* = 11.5, 3.8 Hz, 1H), 2.25 (td, *J* = 10.9, 5.4 Hz, 1H).

*Methyl (R)-4-(2-methoxyethyl)-1-((4-(4-(trifluoromethoxy)phenoxy)phenyl)sulfonyl) piperazine-2-carboxylate* (**7**). **3** (579 mg, 1.03 mmol), TFA (0.79 mL, 10.30 mmol), DCM (10 mL) to give crude mass of 429 mg. Potassium carbonate (386 mg, 2.79 mmol), DMF (5 mL), 2-bromoethyl methyl ether (131 µL, 1.39 mmol) following column elution from 40% to 100% ethyl acetate in hexane to give **7** (270 mg, 0.52 mmol, 50%) as colourless liquid. $^1$H NMR (400 MHz, $CDCl_3$) δ 7.77 (d, *J* = 8.8 Hz, 2H), 7.26 – 7.20 (m, 2H), 7.12 – 7.00 (m, 4H), 4.67 – 4.58 (m, 1H), 3.71 – 3.53 (m, 4H), 3.51 – 3.33 (m, 4H), 3.31 (s, 3H), 2.85 – 2.71 (m, 1H), 2.66 – 2.46 (m, 2H), 2.45 – 2.34 (m, 1H), 2.33 – 2.20 (m, 1H).

*Methyl (R)-4-(2-methoxyethyl)-1-((4-(4-methoxyphenoxy)phenyl)sulfonyl)piperazine-2-carboxylate* (**8**). **4** (400 mg, 0.79 mmol), TFA (0.60 mL, 7.90 mmol), DCM (10 mL) to give crude mass of 235 mg. Potassium carbonate (120 mg, 0.87 mmol), DMF (5 mL), 2-bromoethyl methyl ether (82 µL, 0.87 mmol) following column elution from 40% to 100% ethyl acetate in hexane to give **8** (155 mg, 0.33 mmol, 42%) as pale-yellow liquid. $^1$H NMR (400 MHz, $CDCl_3$) δ 7.73 (d, *J* = 8.8 Hz, 2H), 7.06 – 6.91 (m, 6H), 4.65 – 4.61 (m, 1H), 3.85 (s, 3H), 3.66 – 3.56 (m, 1H), 3.63 (s, 3H), 3.50 – 3.35 (m, 4H), 3.32 (s, 3H), 2.82 – 2.75 (m, 1H), 2.65 – 2.48 (m, 2H), 2.40 (dd, *J* = 11.5, 3.9 Hz, 1H), 2.26 (td, *J* = 11.2, 5.6 Hz, 1H).

*Methyl (R)-4-(2-methoxyethyl)-1-((4-methoxyphenyl)sulfonyl)piperazine-2-carboxylate* (**11**). **5** (675 mg, 2.56 mmol), TFA (1.96 mL, 25.60 mmol), DCM (10 mL) to give crude mass of 456 mg. Potassium carbonate (300 mg, 2.17 mmol), DMF (10 mL), 2-bromoethyl methyl ether (177 µL, 1.88 mmol) following column elution from 40% to 100% ethyl acetate in hexane to give **11** (329 mg, 0.88 mmol, 34%) as colourless liquid. $^1$H NMR (400 MHz, $CDCl_3$) δ 7.72 (d, *J* = 8.8 Hz, 2H), 6.95 (d, *J* = 8.8 Hz, 2H), 4.63 – 4.58 (m, 1H), 3.86 (s, 3H), 3.59 (s, 3H), 3.58 – 3.52 (m, 1H), 3.48 – 3.38 (m, 2H), 3.38 – 3.31 (m, 2H), 3.29 (s, 3H), 2.79 – 2.71 (m, 1H), 2.61 – 2.44 (m, 2H), 2.36 (dd, *J* = 11.5, 3.9 Hz, 1H), 2.22 (td, *J* = 11.5, 3.6 Hz, 1H).

**Methyl (R)-1-((4-methoxyphenyl)sulfonyl)piperazine-2-carboxylate (9).**

In a 50 mL round bottom flask, N-Boc protected **5** (1.60 g, 3.86 mmol) was treated with trifluoroacetic acid (5 mL) in DCM (10 mL) and the reaction mixture was stirred for 16 h. The reaction mixture was concentrated and re-dissolved in DCM. The organic solution was washed carefully with saturated solution of sodium bicarbonate, dried with $MgSO_4$, filtered and concentrated *in-vacuo* to give **9** (1.08 g, 3.44 mmol, 89%) as white crystals. $^1$H NMR (400 MHz, $CDCl_3$) δ 7.74 – 7.68 (d, *J* = 8.8 Hz, 2H), 7.00 – 6.91 (d, *J* = 8.8 Hz, 2H), 4.55 (m, 1H), 3.86 (s, 3H), 3.62 – 3.52 (m, 4H), 3.42 – 3.35 (m, 1H), 3.33 – 3.24 (m, 1H), 3.02 – 2.91 (m, 2H), 2.77 (td, *J* = 12.2, 3.7 Hz, 1H).

**(R)-1-((4-(4-chlorophenoxy)phenyl)sulfonyl)-N-hydroxy-4-(2-methoxyethyl)piperazine-2-carboxamide (15).**

In a 25 mL round bottom flask, methyl ester **4** (36 mg, 0.076 mmol) was dissolved in methanol (1 mL). Aqueous 2 M sodium hydroxide (0.38 mL, 0.76 mmol) solution was added and the reaction was stirred for 1 h. The reaction mixture was concentrated *in-vacuo* and water was added. The aqueous was washed with ethyl acetate (2 times). The aqueous solution was acidified to pH 3-4 with aqueous HCl (2 M) and the reaction mixture was extracted with ethyl acetate (3 times). The pooled organic solution was dried with $MgSO_4$, filtered and concentrated *in-vacuo* to give crude carboxylic acid (28 mg, 0.061 mmol, 80%) and was used without further purification. $^1$H NMR (400 MHz, $CDCl_3$) δ 9.62 (br s, 1H), 7.81 (d, *J* = 8.8 Hz, 2H), 7.35 (d, *J* = 8.8 Hz, 2H), 7.02 - 6.93 (m, 4H), 4.56 – 4.51 (m, 1H), 3.91 – 3.83 (m, 1H), 3.73 – 3.53 (m, 4H), 3.34 – 3.25 (m, 4H), 3.17 – 3.07 (m, 1H), 2.93 – 2.83 (m, 1H), 2.83 - 2.74 (m, 1H), 2.71

– 2.59 (m, 1H). A 25 mL round bottom flask was charge with the carboxylic acid (28 mg, 0.061) starting material, hydroxybenzotriazole (25.0 mg, 0.185 mmol), 1-ethyl-3-(3-dimethylaminopropyl)carbodiimide (0.185 mmol, 35 mg) and THP-protected hydroxylamine (22 mg, 0.185 mmol). Anhydrous DMF (5 mL) and *N*-methylmorpholine (20 µL, 0.185 mmol) were sequentially added and the reaction was stirred overnight. Water was added and the aqueous solution was extracted with ethyl acetate (3 times). The pooled organic solution was dried with $MgSO_4$, filtered and concentrated *in-vacuo*. The crude product was purified by column chromatography following column elution from 50% to 100% ethyl acetate in hexane to give OTHP-amide coupled product (30 mg, 0.055 mmol, 89%) as colourless liquid. $^1$H NMR (400 MHz, $CDCl_3$) δ 7.81 (d, *J* = 9.2 Hz, 2H), 7.39 – 7.28 (m, 6H), 4.70 - 4.50 (m, 2H), 3.90 – 3.80 (m, 1H), 3.80 - 3.70 (m, 1H), 3.55 – 3.42 (m, 2H), 3.36 – 3.27 (m, 2H), 3.29 (s, 3H), 3.03 – 2.93 (m, 1H), 2.84 – 2.62 (m, 2H), 2.61 – 2.45 (m, 2H), 1.79 – 1.45 (m, 7H). In a 25 mL round bottom flask, the resulting OTHP-protected hydroxyl amine (30 mg, 0.054 mmol) was dissolved in 1,4-dioxane (5 mL). 4 N HCl (1 mL in dioxane) was added and the reaction was stirred for 1 h. The reaction mixture was neutralized to pH 7 with saturated solution of sodium bicarbonate. The solution was extracted with ethyl acetate (3 times). The pooled organic solution was dried with $MgSO_4$, filtered and concentrated *in-vacuo*. The crude product was purified by column chromatography following column elution of 10% methanol in DCM to give **15** (151 mg, 0.32 mmol, 52%) as white crystals. $^1$H NMR (400 MHz, $CDCl_3$) δ 7.85 (d, *J* = 9.2 Hz, 2H), 7.35 (d, J = 9.2 Hz, 2H), 7.05 – 6.96 (m, 4H), 4.69 – 4.64 (m, 1H), 3.73 – 3.64 (m, 1H), 3.55 – 3.41 (m, 2H), 3.37 (s, 3H), 3.22 – 3.15 (m, 1H), 3.07 (td, *J* = 12.8, 3.1 Hz, 1H), 2.82 – 2.75 (m, 1H), 2.72 – 2.63 (m, 1H), 2.54 – 2.46 (m, 1H), 2.41 – 2.31 (m, 2H). $^{13}$C NMR (126 MHz, $CDCl_3$) δ 165.43, 161.25, 154.00, 133.62, 130.30, 130.26, 130.13, 121.63, 117.68, 69.21, 59.15, 56.52, 55.49, 52.85, 52.40, 42.62. HMRS (ESI) calculated for $C_{20}H_{24}ClN_3O_6S^+$ $[M+H]^+$ m/z 470.1147, found 470.1148. HPLC purity: 95.0% (RT: 8.59 min).

**General procedure for the conversion of methyl ester to hydroxylamine 16-23 and 30.**

Methyl ester was first dissolved 1,4-dioxane. Potassium cyanate (KOCN) followed by aqueous 50% aqueous hydroxylamine was added and the reaction was stirred for 16 h. The reaction mixture was then concentrated, water and brine were added. The aqueous solution was extracted with ethyl acetate (3 times). The pooled organic solution was dried with $MgSO_4$, filtered and concentrated *in-vacuo*. The crude product was purified by column chromatography. Unless otherwise stated, the purified mass undergoes subsequent semi-preparative HPLC purification to achieve product purity above 95%.

*(R)-N-hydroxy-4-(2-methoxyethyl)-1-((4-(4-(trifluoromethoxy)phenoxy)phenyl)sulfonyl)piperazine-2-carboxamide* (**16**). Methyl ester **7** (165 mg, 0.32 mmol), 1,4-dioxane (8 mL), KOCN (36 mg, 0.44 mmol), 50% aqueous hydroxylamine (8 mL) following column elution from 60% to 100% ethyl acetate in hexane to give **16** (85 mg, 0.16 mmol, 52%) as a colourless liquid. $^1$H NMR (400 MHz, $CDCl_3$) δ 7.87 (d, *J* = 8.9 Hz, 2H), 7.24 (d, *J* = 8.9 Hz, 2H), 7.07 (d, *J* = 9.2, 2H), 7.03 (d, *J* = 9.2 Hz, 2H), 4.69 – 4.65 (m, 1H), 3.74 – 3.65 (m, 1H), 3.57 – 3.41 (m, 2H), 3.36 (s, 3H), 3.30 – 3.22 (m, 1H), 3.18 – 3.12 (m, 1H), 2.86 – 2.80 (m, 1H), 2.73 – 2.67 (m, 1H), 2.59 – 2.52 (m, 1H), 2.41 – 2.34 (m, 2H). $^{13}$C NMR (101 MHz, $CDCl_3$) δ 165.44, 161.16, 153.85, 145.79 (q, *J* = 1.9 Hz), 133.74 (m), 130.29, 123.04, 121.34, 120.36 (q, *J* = 256.84 Hz), 117.86, 68.89, 58.95, 56.40,

55.12, 52.59, 52.25. HRMS (ESI) calculated for $C_{21}H_{25}F_3N_3O_7S^+$ $[M+H]^+$ m/z 520.1360, $C_{42}H_{48}F_6N_6NaO_{14}S_2^+$ $[2M+Na]^+$ m/z 1061.2466, found 520.1374 and 1061.2479. HPLC purity: 96.5% (RT: 7.73 min).

*(R)-N-hydroxy-4-(2-methoxyethyl)-1-((4-(4-methoxyphenoxy)phenyl)sulfonyl)piperazine-2-carboxamide* (**17**). Methyl ester **8** (150 mg, 0.32 mmol), 1,4-dioxane (7 mL), KOCN (37 mg, 0.45 mmol), 50% aqueous hydroxylamine (7 mL) following column elution from 60% to 100% ethyl acetate in hexane to give **17** (112 mg, 0.24 mmol, 75%) as a colourless liquid. $^1$H NMR (400 MHz, $CDCl_3$) δ 7.79 (d, $J$ = 8.8 Hz, 2H), 7.05 – 6.87 (m, 6H), 4.65 – 4.60 (m, 1H), 3.82 (s, 3H), 3.69 – 3.62 (m, 1H), 3.54 – 3.40 (m, 2H), 3.36 (s, 3H), 3.23 – 3.16 (m, 1H), 3.15 – 3.05 (m, 1H), 2.79 – 2.71 (m, 1H), 2.68 – 2.59 (m, 1H), 2.53 – 2.44 (m, 1H), 2.33 – 2.22 (m, 2H). $^{13}$C NMR (101 MHz, $CDCl_3$) δ 165.72, 162.65, 156.97, 148.31, 132.57, 130.05, 121.86, 116.74, 115.29, 69.10, 58.94, 56.44, 55.79, 55.28, 52.49, 52.13, 42.44. HRMS (ESI) calculated for $C_{21}H_{28}N_3O_7S^+$ $[M+H]^+$ m/z 466.1642, found 466.1693. HPLC purity: 97.0% (RT: 9.43 min).

*(R)-N-hydroxy-1-((4-methoxyphenyl)sulfonyl)piperazine-2-carboxamide hydrochloride salt* (**18**). Methyl ester **9** (218 mg, 0.69 mmol), 1,4-dioxane (7 mL), KOCN (79 mg, 0.97 mmol), 50% aqueous hydroxylamine (7 mL). The reaction mixture was concentrated and re-dissolved in ethyl acetate (20 mL). The organic solution was washed with water (2 x 20 mL), dried with $MgSO_4$, filtered and concentrated. The resulting crude was treated with 4 N HCl (5 mL in dioxane) and stirred for 15 minutes. The reaction mixture was concentrated and re-dissolved in methanol (1 mL). Dichloromethane was added drop-wise and the precipitate formed was collected by filtration to give **18** (195 mg, 0.55 mmol, 80%) as a white solid. $^1$H NMR (500 MHz, MeOD) δ 7.83 (d, $J$ = 8.9 Hz, 2H), 7.12 (d, $J$ = 8.9 Hz, 2H), 4.65 - 4.60 (m, 1H), 3.95 – 3.86 (m, 4H), 3.70 – 3.58 (m, 2H), 3.36 - 3.32 (m, 1H), 3.15 - 3.09 (m, 1H), 2.96 (td, $J$ = 12.8, 4.1 Hz, 1H). $^{13}$C NMR (126 MHz, MeOD) δ 165.34, 130.91, 130.87, 115.84, 56.32, 50.44, 45.30, 43.69, 40.60. HRMS (ESI) calculated for $C_{12}H_{18}N_3O_5S^+$ $[M+H]^+$ m/z 316.0962, found 316.0968. HPLC purity: 95.0% (RT: 0.82 min).

*(R)-N-hydroxy-1-((4-methoxyphenyl)sulfonyl)-4-(methylsulfonyl)piperazine-2-carboxamide* (**19**). Methyl ester **10** (412 mg, 1.05 mmol), 1,4-dioxane (10 mL), KOCN (119 mg, 1.47 mmol), 50% aqueous hydroxylamine (10 mL) following column elution from 60% to 100% ethyl acetate in hexane to give **19** (194 mg, 0.49 mmol, 47%) as a white solid. $^1$H NMR (500 MHz, $CDCl_3$) δ 9.46 (br s, 1H), 7.78 (d, $J$ = 8.9 Hz, 2H), 7.02 (d, $J$ = 8.9 Hz, 2H), 4.68 – 4.62 (m, 1H), 4.23 – 4.16 (m, 1H), 3.89 (s, 3H), 3.88 – 3.80 (m, 1H), 3.62 – 3.54 (m, 1H), 3.39 – 3.29 (m, 1H), 2.83 (s, 3H), 2.71 – 2.62 (m, 1H), 2.61 – 2.52 (m, 1H). $^{13}$C NMR (101 MHz, $CDCl_3$) δ 165.57, 163.92, 130.27, 129.65, 115.10, 55.92, 54.31, 44.61, 43.50, 43.08, 37.60. HRMS (ESI) calculated for $C_{13}H_{20}N_3O_7S_2^+$ $[M+H]^+$ m/z 394.0737 and $C_{13}H_{19}N_3NaOS_2^+$ $[M+Na]^+$ m/z 416.0557, found 394.0742 and 416.0558. HPLC purity: 99.5% (RT: 6.55 min).

*(R)-N-hydroxy-4-(2-methoxyethyl)-1-((4-methoxyphenyl)sulfonyl)piperazine-2-carboxamide* (**20**). Methyl ester **11** (153 mg, 0.41 mmol), 1,4-dioxane (7 mL), KOCN (46 mg, 0.57 mmol), 50% aqueous hydroxylamine (7 mL) following column elution from 60% to 100% ethyl acetate in hexane to give **20** (101 mg, 0.27 mmol, 66%) as a colourless liquid. $^1$H NMR (400 MHz, $CDCl_3$) δ 7.81 (d, $J$ = 8.9 Hz, 2H), 6.96 (d, $J$ = 8.9 Hz, 2H), 4.66 – 4.58 (m, 1H), 3.86 (s, 3H), 3.71 – 3.62 (m, 1H), 3.54 – 3.39 (m, 2H), 3.36 (s, 3H), 3.24 – 3.16 (m, 1H), 3.16 – 3.05 (m, 1H), 2.77 – 2.70 (m, 1H), 2.66 – 2.57 (m, 1H), 2.51 – 2.43 (m, 1H), 2.29 – 2.17 (m, 2H). (101

MHz, $CDCl_3$) δ 165.77, 163.18, 131.19, 129.99, 114.27, 69.13, 58.94, 56.52, 55.73, 55.25, 52.39, 52.06, 42.46. HRMS (ESI) calculated for $C_{15}H_{24}N_3O_6S^+$ $[M+H]^+$ m/z 374.1380, found 374.1376. HPLC purity: 97.0% (RT: 4.98 min).

*(R)-N-hydroxy-1,4-bis((4-methoxyphenyl)sulfonyl)piperazine-2-carboxamide* (**21**). Methyl ester **12** (380 mg, 0.78 mmol), 1,4-dioxane (19 mL), KOCN (46 mg, 0.57 mmol), 50% aqueous hydroxylamine (19 mL) following column elution from 60% to 100% ethyl acetate in hexane, and subsequently recrystallised using DCM to give **21** (277 mg, 0.57 mmol, 73%) as a white solid. $^1$H NMR (400 MHz, DMSO) δ 10.77 (br s, 1H), 9.01 (br s, 1H), 7.64 (d, *J* = 8.8 Hz, 2H), 7.47 (d, *J* = 8.8 Hz, 2H), 7.10 (d, *J* = 9.0 Hz, 2H), 6.92 (d, *J* = 9.0 Hz, 2H), 4.46 - 4.44 (m, 1H), 3.88 (s, 3H), 3.84 – 3.71 (m, 5H), 3.57 - 3.46 (m, 1H), 3.32 - 3.25 (m, 1H), 1.85 – 1.78 (m, 1H), 1.71 – 1.61 (m, 1H). $^{13}$C NMR (101 MHz, DMSO) δ 164.16, 162.96, 162.67, 131.07, 129.70, 129.05, 124.87, 114.37, 55.67, 55.58, 53.00, 45.93, 43.81, 42.14. HRMS (ESI) calculated for $C_{19}H_{24}N_3O_8S_2^+$ $[M+H]^+$ m/z 486.0999 and $C_{19}H_{23}N_3NaO_8S_2^+$ $[M+Na]^+$ m/z 508.0819, found 486.1003 and 508.0817. HPLC purity: 96.0% (RT: 8.01 min).

*(R)-4-((3,5-dimethylisoxazol-4-yl)sulfonyl)-N-hydroxy-1-((4-methoxyphenyl)sulfonyl)piperazine-2-carboxamide* (**22**). Methyl ester **13** (555 mg, 1.17 mmol), 1,4-dioxane (28 mL), KOCN (133 mg, 1.64 mmol), 50% aqueous hydroxylamine (28 mL) following column elution from 60% to 100% ethyl acetate in hexane to give **22** (356 mg, 0.75 mmol, 64%) as a colourless liquid. $^1$H NMR (400 MHz, MeOD) δ 7.77 (d, *J* = 9.0 Hz, 2H), 7.03 (d, *J* = 9.0 Hz, 2H), 4.67 - 4.62 (m, 1H), 4.04 – 3.92 (m, 2H), 3.90 (s, 3H), 3.57 – 3.45 (m, 1H), 3.43 - 3.36 (m, 1H), 2.54 (s, 3H), 2.32 – 2.23 (m, 4H), 2.16 – 2.07 (m, 1H). (101 MHz, MeOD) δ 175.94, 166.74, 165.18, 159.17, 132.34, 130.75, 115.65, 113.14, 56.28, 55.11, 45.71, 44.29, 43.52, 12.95, 11.25. HRMS (ESI) calculated for $C_{17}H_{23}N_4O_8S_2^+$ $[M+H]^+$ m/z 475.0952, found 475.0959. HPLC purity: 99.0% (RT: 7.82 min).

*(R)-4-(cyclopropylsulfonyl)-N-hydroxy-1-((4-methoxyphenyl)sulfonyl)piperazine-2-carboxamide* **23** (DC-174). Methyl ester **14** (360 mg, 0.86 mmol), 1,4-dioxane (18 mL), KOCN (98 mg, 1.21 mmol), 50% aqueous hydroxylamine (18 mL) following column elution from 60% to 100% ethyl acetate in hexane to give **23** (DC-174) (338 mg, 0.80 mmol, 94%) as a colourless liquid. $^1$H NMR (500 MHz, MeOD) δ 7.82 (d, *J* = 8.9 Hz, 2H), 7.10 (d, *J* = 8.9 Hz, 2H), 4.57 – 4.54 (m, 1H), 4.06 – 4.01 (m, 1H), 3.88 (s, 3H), 3.87 – 3.83 (m, 1H), 3.62 - 3.48 (m, 2H), 2.86 (dd, *J* = 12.6, 4.3 Hz, 1H), 2.75 – 2.64 (td, *J* = 11.5, 3.0 Hz, 1H), 2.40 – 2.32 (m, 1H), 1.02 – 0.90 (m, 4H). $^{13}$C NMR (126 MHz, MeOD) δ 167.39, 165.04, 132.31, 130.76, 115.70, 56.27, 55.20, 47.67, 45.63, 43.97, 27.01, 4.93, 4.74. HRMS (ESI) calculated for $C_{15}H_{22}N_3O_7S_2^+$ $[M+H]^+$ m/z 420.0894, found 420.0899. HPLC purity: 98.0% (RT: 7.16 min). Elemental Analysis calculated for $C_{15}H_{21}N_3O_7S_2$ requires C, 42.96%, H, 4.99%, N, 9.74%, S, 15.14%. Found C, 42.99%, H, 5.03%, N, 9.70%, S, 15.19%. ($^1$H and $^{13}$C NMR spectra can be found in Supplementary figures 9 and 10)

*(R)-1-((4'-fluoro-[1,1'-biphenyl]-4-yl)sulfonyl)-N-hydroxy-4-(2-methoxyethyl)piperazine-2-carboxamide* (**30**). Methyl ester **27** (166 mg, 0.38 mmol), 1,4-dioxane (8 mL), KOCN (36 mg, 0.44 mmol), 50% aqueous hydroxylamine (8 mL) following column elution from 60% to 100% ethyl acetate in hexane to give **30** (130 mg, 0.30 mmol, 78%) as a colourless liquid. $^1$H NMR (400 MHz, $CDCl_3$) δ 7.95 (d, *J* = 8.4 Hz, 2H), 7.66 (d, *J* = 8.4 Hz, 2H), 7.61 – 7.54 (m, 2H), 7.20 – 7.11 (m, 2H), 4.74 – 4.70 (m, 1H), 3.77 – 3.70 (m, 1H), 3.56 – 3.41 (m, 2H), 3.37 (s, 3H), 3.24 – 3.18 (m, 1H), 3.16 – 3.06 (m, 1H), 2.83 – 2.75 (m, 1H), 2.72 – 2.62 (m, 1H), 2.54

– 2.45 (m, 1H), 2.41 – 2.30 (m, 2H). $^{13}$C NMR (101 MHz, $CDCl_3$) δ 165.60, 163.17 (d, $J$ = 249.5 Hz), 144.75, 138.29, 135.55 (d, $J$ = 3.3 Hz), 129.19 (d, $J$ = 9.1 Hz), 128.46, 127.49, 116.15 (d, $J$ = 21.2 Hz), 69.02, 58.95, 56.39, 55.34, 52.60, 52.16, 42.52. HRMS (ESI) calculated for $C_{20}H_{25}FN_3O_5S^+$ $[M+H]^+$ m/z 438.1493, $C_{20}H_{24}FN_3NaO_5S^+$ $[M+Na]^+$ m/z 460.1313, $C_{40}H_{48}F_2N_6NaO_{10}S_2^+$ $[2M+Na]^+$ m/z 897.2734, found 438.1496, 460.1307 and 897.2734. HPLC purity: 95.5% (RT: 6.77 min).

**(2R)-1-((4-bromophenyl)sulfonyl)-4-(2-methoxyethyl)-N-((tetrahydro-2H-pyran-2-yl)oxy)piperazine-2-carboxamide (26).**

In a 25 mL round bottom flask, methyl ester 2**5** (625 mg, 1.48 mmol) was dissolved in methanol (20 mL). Aqueous 2 M sodium hydroxide (8.50 mL, 17.0 mmol) solution was added and the reaction was stirred for 1 h. The reaction mixture was concentrated *in-vacuo* and water was added. The aqueous was washed with ethyl acetate (2 times). The aqueous solution was acidified to pH 3-4 with aqueous 2 M HCl and the reaction mixture was extracted with ethyl acetate (3 times). The pooled organic solution was dried with $MgSO_4$, filtered and concentrated *in-vacuo* to give a crude carboxylic acid (493 mg, 1.21 mmol, 82%) as white solid. The crude carboxylic acid was used without further purification. $^1$H NMR (400 MHz, $CDCl_3$) δ 7.73 - 7.66 (d, $J$ = 6.5 Hz, 2H), 7.62 – 7.54 (d, 2H), 4.51 – 4.46(m, 1H), 3.83 – 3.76 (m, 1H), 3.74 – 3.49 (m, 4H), 3.32 (s, 3H), 3.27 – 3.19 (m, 1H), 3.08 – 3.00 (m, 1H), 2.88 – 2.79 (m, 1H), 2.78 – 2.71 (m, 1H), 2.66 – 2.56 (m, 1H). A 25 mL round bottom flask was charge with the resulting carboxylic acid (493 mg, 1.21 mmol), hydroxybenzotriazole (490 mg, 3.63 mmol), 1-Ethyl-3-(3-dimethylaminopropyl)carbodiimide (696 mg, 3.63 mmol) and THP-protected hydroxylamine (425 mg, 3.63 mmol). Anhydrous DMF (20 mL) and *N*-methylmorpholine (400µL, 3.63 mmol) were sequentially added and the reaction was stirred overnight. Water was added and the aqueous solution was extracted with ethyl acetate (3 times). The pooled organic solution was dried with $MgSO_4$, filtered and concentrated *in-vacuo*. The crude product was purified by column chromatography following column elution from 50% to 100% ethyl acetate in hexane to give **26** (1.01 g, 1.99 mmol, 51%) as pale-yellow liquid. $^1$H NMR (400 MHz, $CDCl_3$) δ 7.80 – 7.71 (m, 2H), 7.62 (d, $J$ = 8.1 Hz, 2H), 4.90 - 4.85 (m, 1Ha), 4.61 - 4.55 (m, H), 4.33 - 4.26 (m, 1Hb), 3.91 - 3.65 (m, 2H), 3.52 - 3.42 (m, 2H), 3.32 (s, 3H), 3.26 - 3.10 (m, 2H), 2.91 - 2.78 (m, 1H), 2.70 – 2.48 (m, 2H), 2.44 – 2.28 (m, 2H), 1.84 – 1.46 (m, 7H).

**Methyl (R)-1-((4'-fluoro-[1,1'-biphenyl]-4-yl)sulfonyl)-4-(2-methoxyethyl)piperazine-2-carboxylate (27).**

A pressure tube was charged with an aryl-bromide **25**, 1,4-dioxane (4 mL), tetrakis(triphenylphosphine)palladium(0) (20.0 mg, 0.017 mmol), (4-fluorophenyl)boronic acid (133 mg, 0.64 mmol) and aqueous 2 M $K_2CO_3$ (1.65 mL). The reaction mixture was heated at 80°C for 16 h. Water was added and the mixture was extracted with ethyl acetate (3 times). The pooled organic solution was dried with $MgSO_4$, filtered and concentrated *in-vacuo*. The crude product was purified by column chromatography following column elution from 10% to 50% ethyl acetate in hexane to give **27** (173 mg, 0.40 mmol, 69%) as a colourless liquid. $^1$H NMR (400 MHz, $CDCl_3$) δ 7.84 (d, $J$ = 8.8 Hz, 2H), 7.65 (d, $J$ = 8.8 Hz, 2H), 7.60 – 7.54 (m, 2H), 7.20 – 7.12 (m, 2H), 4.68 – 4.63 (m, 1H), 3.69 – 3.62 (m, 1H), 3.57 (s, 3H), 3.46 – 3.35 (m, 4H), 3.29 (s, 3H), 2.82 – 2.75 (m, 1H), 2.62 – 2.46 (m, 2H), 2.43 – 2.36 (m, 1H), 2.27 (td, $J$ = 11.5, 3.6 Hz, 1H).

**(R)-N-hydroxy-4-(2-methoxyethyl)-1-((4-(4-(trifluoromethyl)phenoxy)phenyl)sulfonyl)piperazine-2-carboxamide (28).**

A seal tube was charged with aryl bromide **26** (148 mg, 0.29 mmol), 4-(trifluoromethyl)phenol (57 mg, 0.35 mmol), di-t-BuXphos (13 mg, 0.030 mmol), palladium acetate (7 mg, 0.031 mmol), potassium phosphate (124 mg, 0.58 mmol) in anhydrous toluene (5 mL) and the reaction mixture was stirred at 100°C for 16 hours. The reaction mixture was cooled to room temperature and water (30 mL) was added. The aqueous solution was extracted with ethyl acetate (2 x 50 mL). The pooled organic solution was dried with $MgSO_4$, filtered and concentrated *in-vacuo*. The resulting crude biphenyl material was treated with 4 N HCl (2 mL) and stirred for 1 hour. The reaction mixture was neutralized to pH 7 with saturated solution of sodium bicarbonate. The solution was extracted with ethyl acetate (3 times). The pooled organic solution was dried with $MgSO_4$, filtered and concentrated *in-vacuo*. The crude product was purified by column chromatography with column elution from 50% to 90% ethyl acetate in hexane, followed by preparative HPLC purification to **28** (18 mg, 0.031 mmol, 91%) as a colourless liquid. $^1$H NMR (400 MHz, $CDCl_3$) δ 7.89 (d, *J* = 8.8 Hz, 2H), 7.65 (d, *J* = 8.8 Hz, 2H), 7.13 (d, *J* = 8.4 Hz, 2H), 7.08 (d, *J* = 8.4 Hz, 2H), 4.75 - 4.66 (m, 1H), 3.79 - 3.70 (m, 4H), 3.57 – 3.43 (m, 2H), 3.39 (s, 3H), 3.24 - 3.16 (m, 1H), 3.14 - 3.03 (m, 1H), 2.88 - 2.80 (m, 1H), 2.76 - 2.67 (m, 1H), 2.58 - 2.49 (m, 1H), 2.48 - 2.39 (m, 2H). $^{13}$C NMR (126 MHz, $CDCl_3$) δ 165.07, 160.20, 158.64, 134.44, 130.31, 127.65 (d, *J* = 3.9 Hz), 121.94 (q, *J* = 266.1 Hz), 119.71, 119.69 (q, *J* = 11.0 Hz), 118.75 (q, *J* = 12.9 Hz), 118.68, 68.91, 58.96, 56.20, 55.22, 52.76, 52.18, 42.42. HMRS (ESI) calculated for $C_{21}H_{25}F_3N_3O_6S^+$ $[M+H]^+$ m/z 504.1411, found 504.1413. HPLC purity: 95.0% (RT: 7.66 min).

**(R)-1-((4'-chloro-[1,1'-biphenyl]-4-yl)sulfonyl)-N-hydroxy-4-(2-methoxyethyl)piperazine-2-carboxamide (29).**

A pressure tube was charged with an aryl-bromide **26** (109 mg, 0.215 mmol), 1,4-dioxane (4 mL), tetrakis(triphenylphosphine)palladium(0) (8 mg, 0.0070 mmol), (4-chlorophenyl)boronic acid (37 mg, 0.24 mmol) and aqueous 2 M $K_2CO_3$ (0.6 mL). The reaction mixture was heated at 80°C for 16 h. Water was added and the mixture was extracted with ethyl acetate (3 times). The pooled organic solution was dried with $MgSO_4$, filtered and concentrated *in-vacuo*. The crude product was purified by column chromatography following column elution from 50% to 90% ethyl acetate in hexane to give a biphenyl product (91 mg, 0.17 mmol, 79%) as a colourless liquid. $^1$H NMR (400 MHz, $CDCl_3$) δ 8.00 - 7.92 (m, 2H), 7.68 - 7.64 (d, *J* = 8.4 Hz, 2H), 7.53 - 7.51 (dd, *J* = 8.8, 2.4 Hz, 2H), 7.46 - 7.42 (m, 2H), 4.92 - 4.88 (m, 1Ha), 4.65 - 4.59 (m, 1H), 4.35 - 4.28 (m, 1Hb), 3.90 - 3.70 (m, 2H), 3.52 – 3.43 (m, 2H), 3.32 (s, 3H), 3.29 – 3.12 (m, 2H), 2.91 - 2.78 (m, 1H), 2.71 – 2.48 (m, 2H), 2.45 – 2.27 (m, 2H), 1.82 - 1.65 (m, 2H), 1.59 – 1.41 (m, 5H). The resulting biphenyl material (87 mg, 0.16 mmol) was treated with 4 N HCl (2 mL in 1,4-dioxane) and the reaction mixture was stirred for 1 hour. The reaction mixture was neutralized to pH 7 with saturated solution of sodium bicarbonate. The solution was extracted with ethyl acetate (3 times). The pooled organic solution was dried with $MgSO_4$, filtered and concentrated *in-vacuo*. The crude product was purified by column chromatography following column elution of 10% methanol in DCM and subsequent semipreparative HPLC purification to give **29** (60 mg, 0.13 mmol, 81%) as colourless liquid. $^1$H NMR (400 MHz, $CDCl_3$) δ 8.00 (s, 1H), 7.95 (d, *J* = 8.3 Hz, 2H), 7.66 (d, *J* = 8.3 Hz, 2H), 7.54 (d, *J* = 8.4 Hz, 2H), 7.44 (d, *J* = 8.4 Hz, 2H), 4.76 - 4.70 (m, 1H), 3.81 – 3.72 (m, 1H), 3.56 – 3.41 (m, 2H), 3.37 (s, 3H), 3.26 – 3.18 (m, 1H), 3.18 – 3.07 (m, 1H), 2.87 – 2.78 (m, 1H), 2.74 – 2.64 (m,

1H), 2.57 – 2.47 (m, 1H), 2.46 – 2.35 (m, 2H). $^{13}$C NMR (126 MHz, $CDCl_3$) δ 165.20, 151.01, 144.52, 137.91, 134.87, 129.37, 128.73, 128.50, 127.48, 68.89, 58.97, 56.27, 55.23, 52.69, 52.16, 42.43. HRMS (ESI) calculated for $C_{20}H_{25}ClN_3O_5S^+$ $[M+H]^+$ m/z 454.1198, $C_{20}H_{24}ClN_3NaO_5S^+$ $[M+Na]^+$ m/z 476.1017, $C_{40}H_{48}Cl_2N_6NaO_{10}S_2^+$ [2M+Na]+ m/z 929.2143, found 454.1196, 476.1010 and 929.2136. HPLC purity: 96.0% (RT: 7.27 min).

*Supplementary References*


1 Lingott, T., Schleberger, C., Gutiérrez, J. M. & Merfort, I. High-Resolution Crystal Structure of the Snake Venom Metalloproteinase BaP1 Complexed with a Peptidomimetic: Insight into Inhibitor Binding. *Biochemistry* **48**, 6166-6174 (2009). https://doi.org:10.1021/bi9002315

2 Zhang, D. *et al.* Structural interaction of natural and synthetic inhibitors with the venom metalloproteinase, atrolysin C (form d). *Proceedings of the National Academy of Sciences* **91**, 8447-8451 (1994). https://doi.org:10.1073/pnas.91.18.8447

3 Misson Mindrebo, L. E. *et al.* Importance of the Cysteine-Rich Domain of Snake Venom Prothrombin Activators: Insights Gained from Synthetic Neutralizing Antibodies. *Toxins (Basel)* **16** (2024). https://doi.org:10.3390/toxins16080361

4 Takeda, S., Igarashi, T. & Mori, H. Crystal structure of RVV-X: An example of evolutionary gain of specificity by ADAM proteinases. *FEBS Letters* **581**, 5859-5864 (2007). https://doi.org/10.1016/j.febslet.2007.11.062

5 Sievers, F. *et al.* Fast, scalable generation of high-quality protein multiple sequence alignments using Clustal Omega. *Molecular Systems Biology* **7**, 539 (2011). https://doi.org/10.1038/msb.2011.75

6 Waterhouse, A. *et al.* SWISS-MODEL: homology modelling of protein structures and complexes. *Nucleic Acids Research* **46**, W296-W303 (2018). https://doi.org:10.1093/nar/gky427

7 Studer, G. *et al.* QMEANDisCo—distance constraints applied on model quality estimation. *Bioinformatics* **36**, 2647-2647 (2020). https://doi.org:10.1093/bioinformatics/btaa058

8 Jones, G., Willett, P., Glen, R. C., Leach, A. R. & Taylor, R. Development and validation of a genetic algorithm for flexible docking11Edited by F. E. Cohen. *Journal of Molecular Biology* **267**, 727-748 (1997). https://doi.org/10.1006/jmbi.1996.0897

9 Lovejoy, B. *et al.* Crystal structures of MMP-1 and -13 reveal the structural basis for selectivity of collagenase inhibitors. *Nature Structural Biology* **6**, 217-221 (1999). https://doi.org:10.1038/6657

10 Choi, J. Y. & Chung, E. Molecular Dynamics Simulations of Matrix Metalloproteinase 13 and the Analysis of the Specificity Loop and the S1′−Site. *International Journal of Molecular Sciences* **24** (2023). https://doi: 10.3390/ijms241310577

11 Campbell, A. J., Lamb, M. L. & Joseph-McCarthy, D. Ensemble-Based Docking Using Biased Molecular Dynamics. *Journal of Chemical Information and Modeling* **54**, 2127-2138 (2014). https://doi.org:10.1021/ci400729j

12 Lovejoy, B. *et al.* Structure of the Catalytic Domain of Fibroblast Collagenase Complexed with an Inhibitor. *Science* **263**, 375-377 (1994). https://doi.org:10.1126/science.8278810

13 Lindorff-Larsen, K. *et al.* Improved side-chain torsion potentials for the Amber ff99SB protein force field. *Proteins: Structure, Function, and Bioinformatics* **78**, 1950-1958 (2010). https://doi.org/10.1002/prot.22711

14 Bussi, G., Donadio, D. & Parrinello, M. Canonical sampling through velocity rescaling. *The Journal of Chemical Physics* **126**, 014101 (2007). https://doi.org:10.1063/1.2408420

15 Parrinello, M. & Rahman, A. Polymorphic transitions in single crystals: A new molecular dynamics method. *Journal of Applied Physics* **52**, 7182-7190 (1981). https://doi.org:10.1063/1.328693

16 McGibbon, Robert T. *et al.* MDTraj: A Modern Open Library for the Analysis of Molecular Dynamics Trajectories. *Biophysical Journal* **109**, 1528-1532 (2015). https://doi.org/10.1016/j.bpj.2015.08.015

17 Pedregosa, F. *et al.* Scikit-learn: Machine Learning in Python. *J. Mach. Learn. Res.* **12**, 2825–2830 (2011).

18 Badawi, M. M. *et al.* Immunoinformatics Predication and in silico Modeling of Epitope-Based Peptide Vaccine Against virulent Newcastle Disease Viruses. *American Journal of Infectious Diseases and Microbiology* **4**, 61-71 (2016). https://doi.org:10.12691/ajidm-4-3-3
